# Ether linkages in phospholipids provide modular control of membrane mechanics

**DOI:** 10.64898/2026.05.20.723252

**Authors:** Jacob R. Winnikoff, Sasiri J. Vargas-Urbano, Daniel Milshteyn, Diego L. Velasco-González, Petr Shendrik, Elida Kocharian, Kelly N. Isbell, Alexander J. Sodt, Raya Sorkin, Peter R. Girguis, Edward Lyman, Itay Budin

## Abstract

The structures of phospholipid headgroups and chains are well-established determinants of membrane elastic properties, but functions for different chemistries that join these moieties together are poorly understood. While common phospholipids feature ester linkages, alkyl ether- and plasmenyl-linked species emerged in prokaryotes, are highly abundant in metazoans, and have been implicated in neurodegeneration and aging. Multiple pathways for ether lipid synthesis evolved convergently, suggesting conserved functions for ether linkage chemistry in the structure of cell membranes. Here we combine experiments and molecular simulations to show that backbone linkage chemistry provides modular control of membrane mechanics and topology through a set of discrete drivers. Alkyl linkages provide the first of these by additively promoting negative intrinsic curvature, which destabilizes bilayers. Ethers also decouple membrane stiffness from viscosity, softening membranes while maintaining packing in the hydrophobic core. The plasmenyl C=C bond represents a second, distinct driver that stabilizes the inverted hexagonal phase by relieving interstitial packing frustration. These results explain the fusogenicity of ether lipids, show how they regulate membrane topology through multiple physical mechanisms, and provide a rationale for convergent evolution of the complex plasmenyl linkage moiety.

**Significance statement:** The structure and dynamics of cell membranes can be sensitive to small chemical changes in their phospholipid building blocks. We describe how phospholipid backbone linkage chemistry controls membrane mechanical properties: ether linkages facilitate membrane dynamics by imposing a more conical molecular profile and by decoupling bending stiffness from chain ordering. Plasmalogen lipids, in which the ether linkage is modified with a vicinal double bond, further promote non-lamellar topologies through relief of interstitial packing frustration. These biophysical features suggest a basis for the repeated emergence of ether phospholipids in evolution and their observed functions in membrane trafficking. The thermodynamic and structural bases of plasmalogen function are especially notable as these lipids have been increasingly implicated in neurodegenerative and cardiovascular disease.

## Introduction

Phosphoglycerolipids, the fundamental building blocks of cell membranes, contain three structural modules: a phosphate-bearing headgroup, a glycerol backbone, and one or more hydrocarbon (radyl) chains. In cells, both headgroups and chains show a wide range of chemical variations, which modulate the physicochemical properties of membranes. For example, charge of headgroups determines surface electrostatics, while the length and unsaturation of chains are dominant factors in the thickness and fluidity of the membrane. A combination of headgroup size, interactions, and chain structure determines a lipid’s molecular shape, which influences whether it forms flat bilayers (lamellar phases) or highly curved non-bilayer structures, like micelles or inverted phases. These distinct membrane properties are often coupled. Among zwitterionic phospholipids, the headgroup primary ammonium in phosphatidylethanolamine (PE) reduces membrane fluidity compared to the quaternary ammonium in phosphatidylcholine (PC) due to its increased hydrogen bonding, but also favors non-lamellar phases due to its smaller effective size. When incorporated into bilayers, high-curvature lipids like PE impart packing frustration – a curvature elastic strain – that is important for membrane trafficking (1, 2) and membrane protein function (3–5). In vitro, processes like SNARE-mediated membrane fusion require high-curvature species like PE (6), which can comprise the majority of lipids in some cell membranes (7).

Though headgroups and radyl chains are recognized determinants of membrane properties, the functional basis for structural differences in the backbone that connects them is less well understood. Typically, eukaryotic and bacterial phospholipids feature glycerol-phosphate backbones that are acylated with fatty acids via ester linkages. However, ether linkages represent an alternative backbone chemistry in eukaryotes and bacteria, and are the dominant form in archaeal membranes. The three most common ether-linked backbones are monoalkyl (2-acyl), dialkyl, and plasmenyl, the latter with an α-alkenyl (“vinyl”) ether. In some cases, these backbones appear to be selected for their chemical properties that are advantageous in certain environments. For example, the hydrolytic resistance of alkyl ethers is important in hot, acidic environments, such as geothermal springs and deep sea vents (8, 9). In contrast, the heightened reactivity of plasmenyl linkage with reactive oxygen species allows it to serve as a bacterial photooxidation sensor (10), and could underlie proposed roles as a cellular antioxidant (11) or a ferroptotic agent (12) in mammals.

Beyond their chemical stability or reactivity, ether linkages could have strong effects on membrane phase properties and mechanics. In vitro, ether-linked PEs form the inverted hexagonal (H_II_) phase – a non-bilayer aggregate in which headgroups face inward and chains outward – over a larger temperature range than ester-linked PEs (13–17). Plasmalogens purified from cells have been observed to enhance vesicle fusion reactions (18), which proceed through an intermediate with non-bilayer topology. In cells, loss of ether lipid metabolism has been observed to cause defects in an ever-expanding range of membrane trafficking and remodeling processes, including endocytosis (19), caveolae formation (20), cell fusion (21), and lysosomal exocytosis (22). Both trafficking and oxidative-stress functions of ether lipids could be relevant for physiological conditions characterized by their loss, including neurodegeneration (23, 24), aging (25), cardiovascular disease (26), and inherited peroxisomal diseases (27).

Because alkyl and plasmenyl linkages arose convergently across bacteria and eukaryotes yet are consistently concentrated under the “non-bilayer” PE headgroup, we reasoned that backbone linkage chemistry confers fundamental functions related to membrane mechanics and topology. We address this hypothesis by systematic comparison of phospholipids that differ only in backbone linkage chemistry. Using a range of experimental and computational approaches, we report how backbone linkage chemistry acts on membrane mechanics through multiple, discrete mechanisms. Alkyl ether substitution drives one–simultaneously increasing curvature elastic strain in bilayers and softening them; the plasmenyl linkage adds a second – lowering the interstitial packing frustration associated with inverted phase formation. In agreement with previous reports on other ether/ester-lipid comparisons (28, 29), we find that ether linkages have relatively little effect on membrane fluidity or ordered phase transitions. Together, these findings elucidate how ether linkages tune membrane mechanics in a manner that promotes membrane trafficking.

## Results

### Ether linkages evolved repeatedly and are enriched in ethanolamine phospholipids

Ether lipids are widely synthesized across the tree of life through distinct metabolic pathways. In bacteria, alkyl and plasmenyl linkages are synthesized by distinct anaerobic and aerobic enzyme pathways whose distribution across dispersed taxa reflects repeated horizontal gene transfer (Fig. 1A; SI Methods). Eukaryotes acquired only the aerobic, peroxisomal pathway (30, 31), which was subsequently lost in plants and most fungi but regained in one fungal lineage (31) (Fig. 1B). The distribution of two convergently evolved pathways across disparate taxa, which reside across a diversity of habitats, argues that ether linkages did not evolve strictly as responses to a single chemical or physical environment. Notably, monoalkyl ethers are found in mesophilic bacteria (32, 33) and ether:ester ratio decreases with rising temperature in some thermophiles (34), suggesting an adaptive role other than hydrolysis resistance. Similarly, one of the two primary pathways for plasmalogen synthesis evolved in an anaerobic context (35, 36), where the lipids’ radical scavenging capacity is unlikely to have physiological relevance.

**Figure 1:**
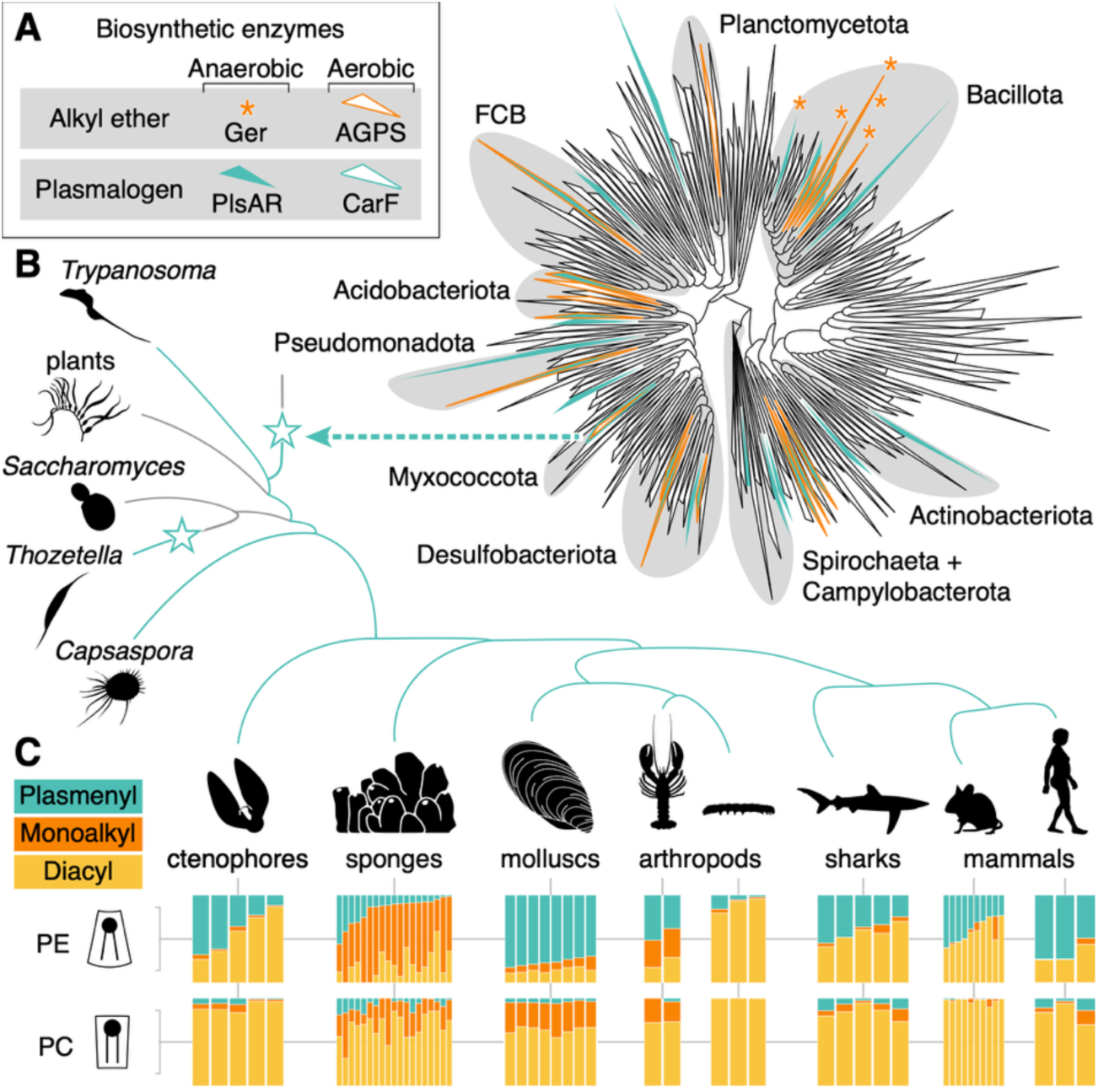
Diversity, biosynthesis and evolution of backbone chemistry in phospholipids. (**A**) Ether phospholipid production is distributed across bacteria. Colored clades represent classes harboring putative orthologs of ether lipid biosynthetic enzymes. For Ger1, only verified ether lipid producers are marked. Select phyla are shaded and labeled. (**B**) Aerobic plasmalogen synthesis appears to have evolved once in Myxococcales, from which CarF/PEDS1 was transferred to early eukaryotes. (**C**) Plasmalogens are present in most crown-group eukaryotes and are concentrated under the PE headgroup. Production of plasmalogens by stem-group eukaryotes, represented in the schematic tree by *Trypanosoma*, suggests that the aerobic synthesis pathway was acquired early. Plasmalogens are absent from plants and most fungi, but were regained by at least one fungal clade. Most later-diverging eukaryotes (e.g. *Capsaspora*) make plasmalogens, and backbone compositions are shown here under PE and PC headgroups in diverse metazoa (5 ctenophores, 19 sponges, 8 bivalve molluscs, 2 motor nerves from the lobster *Homarus gammarus*, 3 larval stages of the tobacco budworm, white muscle from 5 sharks, 9 mouse tissues, and human brain, adipose, and cardiac tissue.) Ether lipid content varies by species, tissue, and life stage, but is consistently biased toward the PE headgroup.

Compiling lipidomic data collected by us and others across metazoans (37–46) (Table S1), we also noted a distinct headgroup bias in ether lipid metabolism (Fig. 1C). Most lineages showed a distinct bias for the plasmenyl vs. monoalkyl linkage in ethanolamine vs. choline phospholipids. Plasmalogen fraction and total ether:ester ratio were found to be consistently higher under non-bilayer PE than under bilayer-forming PC. In some mammalian tissues, PE lipids are dominated by plasmalogens, making plasmenyl PE the second most abundant phospholipid class in several sampled tissues in mice and humans. Because non-bilayer PE lipids exhibit negative intrinsic curvature and are required for topological remodeling of membranes, these patterns are consistent with the notion that ether linkages might modulate related mechanical properties.

### Ether linkages independently promote membrane fusion and inverted phase formation

Membranes must break apart (fission) and come together (fusion) for most forms of cell division (47), as well as intracellular trafficking (48–50). A capacity for biologically derived plasmalogens to promote membrane fusion was first noted thirty years ago (40, 41), but the lack of pure, synthetic ether lipids for systematic comparison limited further dissection of this function. It remains unresolved whether plasmalogens’ fusogenicity is due primarily to the ether linkage, the vicinal C=C bond, or an emergent synergy of the two. With such reagents now available (Fig. 2A), we employed a simple membrane fusion reaction in large unilamellar vesicles (LUVs) containing bilayer (PC) and non-bilayer (PE) zwitterionic lipids, and phosphatidylserine (PS) that interacts with calcium to trigger the reaction (Fig. 2B). In the assay, lipid mixing that occurs during vesicle hemifusion is monitored via FRET dequenching. The unilamellarity and size of ester- and ether-linked PE liposomes was confirmed with a dithionite quenching assay (Fig. S1A-C) and dynamic light scattering (DLS) (Fig. S1D).

**Figure 2:**
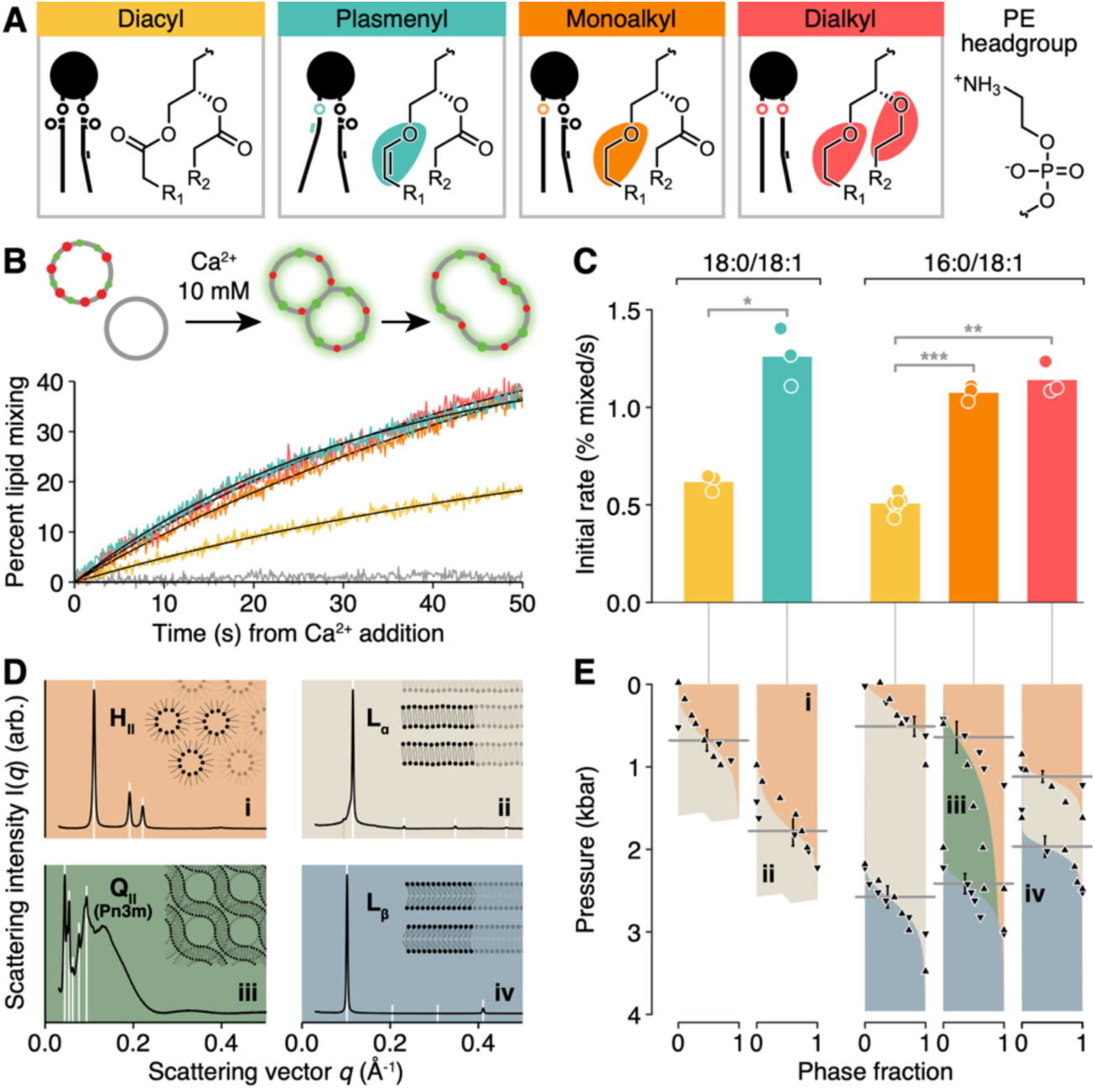
Ether backbone linkages drive large, correlated enhancements of lipid mixing and inverted phase stability. (**A**) Glycerol backbone types considered in this study. Monoalkyl and plasmenyl backbones feature ether linkages at the *sn*-1 position only. In the experiments shown here, all backbones are paired with the PE headgroup shown at right. (**B**) Calcium initiates hemifusion of PS-containing vesicles, detectable as an increase in FRET-donor emission. (**C**) Ether backbone linkages enhance the initial lipid mixing rate. Asterisks indicate significance level; for comparisons indicated by brackets P=0.012, 4.2×10^-3^, 4.3×10^-5^. (**D**) Mesophases formed by PEs and representative SAXS profiles. White lines indicate characteristic peaks; Roman numerals refer to points in (E) where profiles were acquired. The small low-angle peak in (ii) is residual H_II_. Phase cartoons are schematic, not to scale. (**E**) Barotropic phase diagrams measured at 75°C. The plasmenyl linkage drives a large increase in the inversion pressure. Alkyl ether linkages progressively narrow the region between H_II_ and L_β_, with the monoalkyl lipid forming Q_II_ in lieu of L_α_. Horizontal lines mark inflection points of phase transition; error bars are ±SEM.

Using the lipid mixing assay, we compared a series of five PE lipids encompassing the linkage chemistries shown in Fig. 2A. Two diacyl species were used to match the commercially available chain configurations for alkyl and plasmenyl species; they also enabled us to assess the effect of sn-1 chain length. The replacement of diacyl with plasmenyl PE enhanced initial lipid mixing rate, as previously shown (41, 42). However, substitution of monoalkyl PE recapitulated most of the effect of the plasmenyl PE on lipid mixing, relative to the chain-matched diacyl species. Substitution of dialkyl PE, which features an ether linkage at both positions, had no further effect (P = 0.31; Fig. 2C). Across the series, the largest difference in lipid mixing rate was attributable to the replacement of the sn-1 ester with an ether.

Membrane fusion proceeds through a hemifusion “neck” intermediate resembling the non-lamellar H_II_ phase (51, 52). In agreement with this model, it has been noted that the capacity of plasmenyl PE to enhance fusion correlates with its propensity to form H_II_ (53). To test whether the correlation extends to monoalkyl and dialkyl PEs, we constructed barotropic phase diagrams for the same PE series used in lipid mixing experiments. Driving phase transitions with hydrostatic pressure demands more specialized equipment than a temperature scan but perturbs the molecular volume of lipids more selectively, minimizing concomitant changes in chain entropy and interface hydration (54), while better maintaining integrity of the chemically labile plasmenyl linkage. Phases were identified by their distinct SAXS profiles (Fig. 2D) and hydrostatic pressure series were collected in excess pure water at 75°C, a temperature that places all four pictured phases in an accessible pressure range (Fig. 2E). In accordance with our previous results (46), the plasmenyl linkage considerably expanded the range of H_II_ stability (*P_H_*_II→*L*_*_α_* increase of 1097±221 bar, *P*=7.0×10^-7^; two-tailed Wald *z* test). The dialkyl backbone also expanded the H_II_ region, but by a smaller margin (*P_H_*_II→*L*_*_α_* increase of 608±131 bar, *P*=1.2×10^-5^). The monoalkyl PE failed to form L_α_ at any pressure, instead transitioning from H_II_ to an inverted cubic (Q_II_) phase at a pressure similar to *P_H_*_II→*L*_*_α_* of the diacyl analog (*P*=0.56).

Pressure, like low temperature, can drive bilayers from the fluid phase (L_α_) into the gel phase (L_β_) through the ordering of lipid chains. We observed that the dialkyl linkage decreased the L_α_→L_β_ transition pressure compared to the diacyl PE (*P*=8.5×10^-4^), indicating that ether linkages increase chain packing in lamellar phases. The gel transition was not traversed in plasmenyl PE in order to shorten the experiment and protect this chemically delicate sample. Based on these phase diagrams, we concluded that ether linkages destabilize the L_α_ phase in PE lipids, both increasing P_HII→Lα_ and reducing *P_Lα_*_→*L*_*_β_*. This effect contrasts with that of chain unsaturations, which also promote H_II_ but disfavor L_β_ (52, 57, 58).

### Ether linkages contribute to intrinsic curvature in an additive, headgroup-independent manner

One determinant of a phospholipid’s H_II_ propensity is its shape in profile, conventionally described via intrinsic curvature – the geometric curvature adopted by a mechanically relaxed monolayer of the lipid. Lipid intrinsic curvature influences the stored elastic energy of monolayers in the L_α_, H_II_, and hemifusion structures that they form via the monolayer form of the Helfrich Hamiltonian:

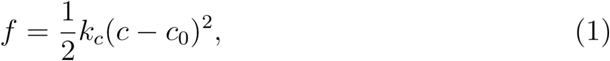

where *f* is elastic energy per unit area, *k_c_* is the curvature (i.e. bending) modulus of the monolayer, *c* is total monolayer curvature, and *c*_0_ is spontaneous monolayer curvature (59). Spontaneous curvature is an elastic parameter that characterizes the residual bending torque in a flat monolayer. In single-component and ideally mixed binary lipid systems, the values of spontaneous and intrinsic curvature are expected to be similar (60, 61). The curvature (reciprocal radius) of an unstressed monolayer can be inferred from the repeat spacing of H_II_ tubules in the presence of excess water and a “filler” hydrocarbon such as 9*Z*-tricosene (TS) to relieve interstitial packing frustration (62). We employed this approach to measure *c*_0_ across the PE series in Figure 2, plus PC analogs for four of those species (Fig. 3A). Since *c*_0_ of most PE lipids is negative by convention, we report −*c*_0_ here, so that larger values correspond to more highly curved lipids.

**Figure 3:**
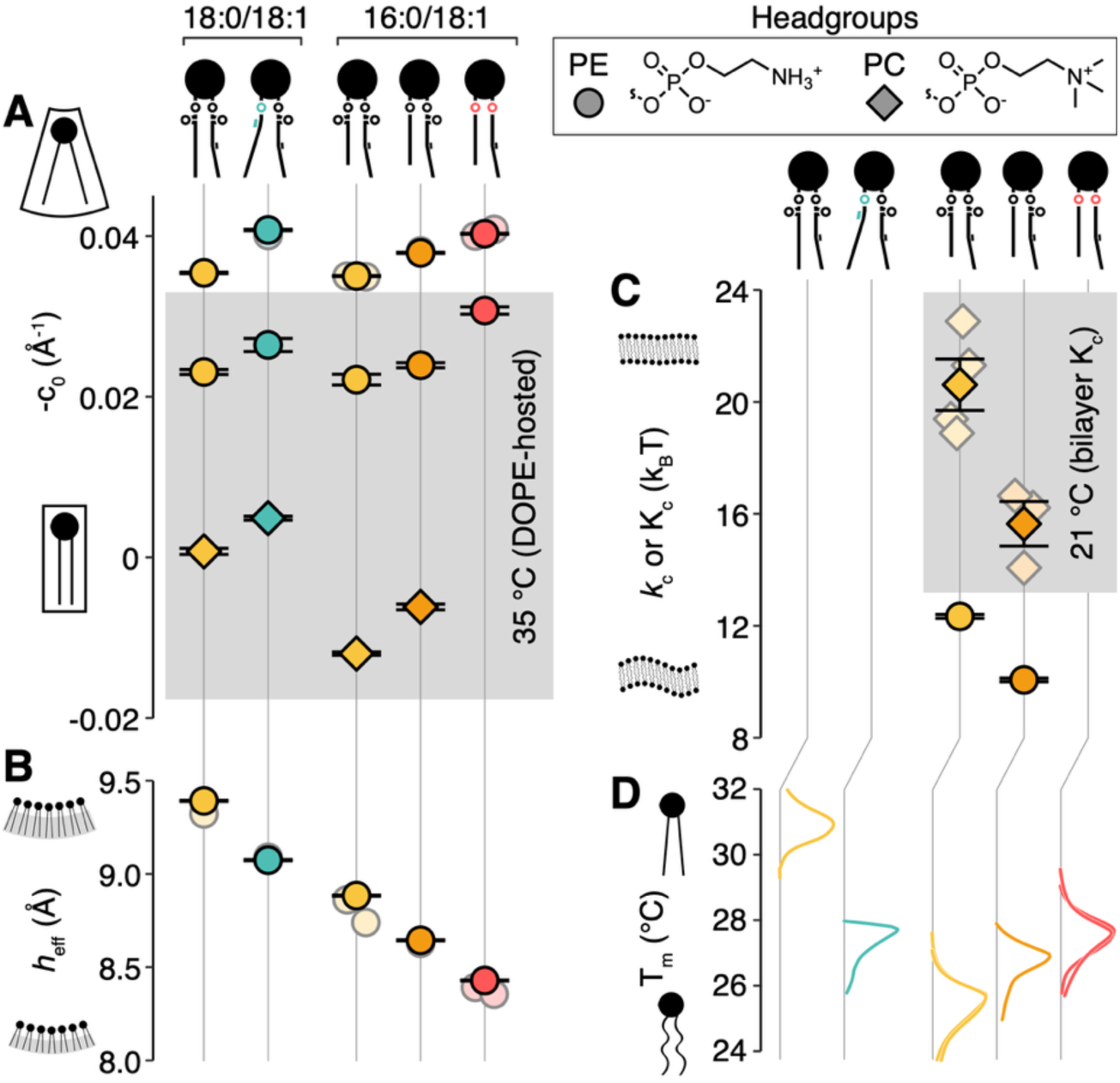
Effect of backbone chemistry on lipid-intrinsic biophysical properties. All values are reported at 75°C and 1 bar unless otherwise noted. Within chain-structure groups, all comparable values differ significantly from each other (*P*_adj_<0.05, pairwise z-tests with Holm-Sidak correction; Table S5A). (**A**) Ether linkages enhance negative intrinsic curvature (−*c*_0_) as measured by SAXS. Increased sn-1 chain length enhances −*c*_0_ slightly. Both effects hold for PE and PC headgroups. To estimate −*c*_0_ at 35°C (shaded area), guest lipids were hosted at 20% (PEs) or 10% (PCs) in DOPE. For pure samples at 75°C, error bars are ±SEM of a jointly fit robust linear model (Table S4A,B); locally fit 1-bar values are shown in the background. Error bars for 35°C are ±SEM from linear regressions vs. pressure. (**B**) Ether linkages yield thinner monolayers. Monolayer effective hydrocarbon thickness (ℎ_eff_) was calculated from H_II_ lattice spacing. Data are fitted and presented similarly to the 75°C data in (A) (Table S4A,C). (**C**) Sn-1 alkyl linkage softens PE monolayers and PC bilayers. Monolayer bending modulus *k_c_* was directly measured for diacyl and monoalkyl POPE using osmotic stress. Bilayer bending modulus *K_c_* (shaded area) was measured in POPC GUVs. Error bars are ±SEM of fitted models (see Methods). (**D**) Ether linkages have consistent but small impacts on chain melting temperature (*T_m_*) as measured by DSC – normalized heat flow traces are shown whose peaks reflect the *T_m_*. Effects of plasmenyl and dialkyl backbones are opposite in direction but similar in magnitude. Both are small; a C_2_ difference in *sn*-1 chain length has about twice the impact of backbone chemistry. Three replicate samples are superimposed.

Using standard geometric relations and partial molar volume corrections (Methods, Tables S2-3), we calculated −*c*_0_ at the pivotal plane (the plane within the monolayer whose cross-sectional area is unchanged by bending) in pure single-lipid systems at 75°C and in “hosted” systems, where the target “guest” lipid is doped into a di-oleoyl PE (DOPE) matrix, at 35°C. As expected, the 40°C temperature difference had a large effect on −*c*_0_, but the qualitative trend with respect to backbone chemistry was consistent across the pure and hosted samples. The −*c*_0_values recovered for diacyl POPE and POPC at 35 °C were ∼0.01 Å^-1^ smaller than those reported in literature (53) at the neutral plane (the plane within the monolayer where bending is decoupled from stretching). We attribute this discrepancy to the difference in reference plane and to error in molecular volume parameters used in our analysis (see Methods, Table S2.)

Our analysis showed that the plasmenyl backbone increased −*c*_0_, consistent with our previous measurements (46) but in contrast to earlier analyses that lacked access to synthetic-defined lipids (16). Our comparison attributed this effect to the ether linkage rather than the vicinal C=C bond, with the dialkyl backbone driving a −*c*_0_ comparable to that of plasmenyl PE. Substituting PC for PE headgroups lowered −*c*_0_ substantially, but monoalkyl and plasmenyl backbones both increased −*c*_0_ by comparable amounts. Taken together, our measurements showed that ether lipids are characterized by more negative intrinsic curvature than their diacyl analogs, that this effect is additive with each ether linkage, and that it is independent of the headgroup.

### Ether linkages decouple flexibility from fluidity

The bending stiffness of lipid monolayers and bilayers is determined by their thickness (65), which generally correlates with chain ordering and bulk viscosity. Stiffness, described by a modulus (*k_c_* for a monolayer; *K_c_* for a bilayer) enters linearly into stored elastic energy alongside the quadratic contribution of *c*_0_ (Eq. 1). We sought to understand the effect of ether linkages on stiffness and whether this effect was bound to its typical covariates. SAXS analysis yielded the mechanical effective hydrocarbon thickness ℎ_eff_of the H_II_ monolayer, defined as the distance between the pivotal plane and chain termini. We found that ether linkages thinned the monolayer in an additive manner resembling the pattern of *c*_0_ values (Fig. 3B). We therefore expected them to soften both monolayers and bilayers: for a symmetric bilayer with uncoupled leaflets, *K_c_* ≈ 2*k_c_*.

Bending moduli (Fig. 3C) were measured directly in PE monolayers using an osmotic stress technique (63) and in PC bilayers using optical tweezer tether-pulling and micropipette aspiration. Different observables (*k_c_* at 75 °C vs. *K_c_* at 21 °C) and techniques were dictated by the non-lamellar and lamellar tendencies of the two headgroups. The host-guest strategy is not viable for measuring *k_c_* due to diffusional softening in mixed systems (13, 66, 67). In both experiments, substitution of the monoalkyl for the diacyl backbone caused softening. In POPE, *k_c_* was reduced by 18.5±0.8%; in POPC, *K_c_* was reduced by 24.1±5.1% (Table S5A). The similarity in effect sizes on *k_c_* and *K_c_* for monoalkyl PE and PC, respectively, suggests that headgroup structure and inter-leaflet coupling do not have large effects on linkage-mediated softening. In PE, the relationship between *k_c_* and ℎ_eff_ was qualitatively consistent with elastic plate theory (55), however the measured softening somewhat exceeded the predicted quadratic scaling (Fig. S2).

In the literature, there has been debate over whether ether linkages, especially plasmalogens, increase or decrease chain packing (68). To resolve this, we measured gel-to-fluid transition temperatures (*T_m_*) of the PE series by microscale differential scanning calorimetry (DSC; Fig. 3D). Monoalkyl PE exhibited *T_m_* 1.2 °C higher than its diacyl analog, and this effect was augmented (+0.8 °C) in dialkyl PE. Plasmenyl PE featured a *T_m_* 3.2 °C lower than its diacyl analog. This *T_m_* drop is consistent with packing interference by the O-vicinal C=C bond, but is dramatically smaller than the >37 °C *T_m_* difference between 18:0/18:1 diacyl PE and 18:1/18:1 diacyl PE caused by addition of the *n*-9 C=C bond (69). A C_2_ difference in *sn*-1 acyl chain length had about twice the maximal impact of backbone chemistry. The same patterns were observed in *P_Lα_*_→*L*_*_β_* measured by SAXS (Fig. 2E), and differences in both the thermotropic and barotropic phase transitions were corroborated by solvatochromic fluorescence measurements (Fig. S4).

These results indicate that the typical relationship between membrane thickness, stiffness, and *T_m_* across lipids is inverted by backbone chemistry: thinner, more flexible ether-linked lipids are surprisingly more gel-prone. Overall, the effects of backbone type on *T_m_* were consistent but small, implying that large fractions of ether lipid can be incorporated into membranes with modest changes to bulk chain packing. This contrasts with other phospholipid features that cells can use to soften membranes, such as shorter and more unsaturated chains, which also reduce chain ordering and alter associated properties like permeability. Given the pronounced effect of ether linkages on flexibility and their minimal impact on fluidity, we propose that ether linkages enable organisms to decouple these two physical properties.

### Simulations trace lipid curvature and membrane flexibility effects to reduced backbone sterics

To identify conformational and dynamical differences underlying the properties of ester- and ether-linked phospholipids, we ran atomistic molecular dynamics (MD) simulations of single-component bilayers at 35°C and 1 bar, consisting of each of the five PE species used in experiments (Fig. 4A). We obtained *c*_0_ and *k_c_* from the simulated bilayers by lateral-pressure (Fig. S5) and transverse curvature bias (TCB) analysis (58) (Methods). For technical reasons, −*c*_0_ is reported at the pivotal plane from experiments and at the neutral plane from simulations, making these values comparable in trend but not in absolute magnitude (see Methods, Table S6A). We found that in simulations, as in experiments, substitution of alkyl for acyl linkages increases −*c*_0_ (Fig. 4B, Table S5B) and decreases *k_c_* (Fig. 4C; Table S5B). However, the plasmenyl systems showed a reduced −*c*_0_ inconsistent with experiment (Table S6A). Because the CHARMM ether lipid force field was parameterized for alkyl linkages (69) and does not account for the unusual electrostatics of the vicinal C=C, we surmised that the plasmenyl force field requires further refinement and focused further MD analyses exclusively on diacyl, monoalkyl, and dialkyl PE.

**Figure 4:**
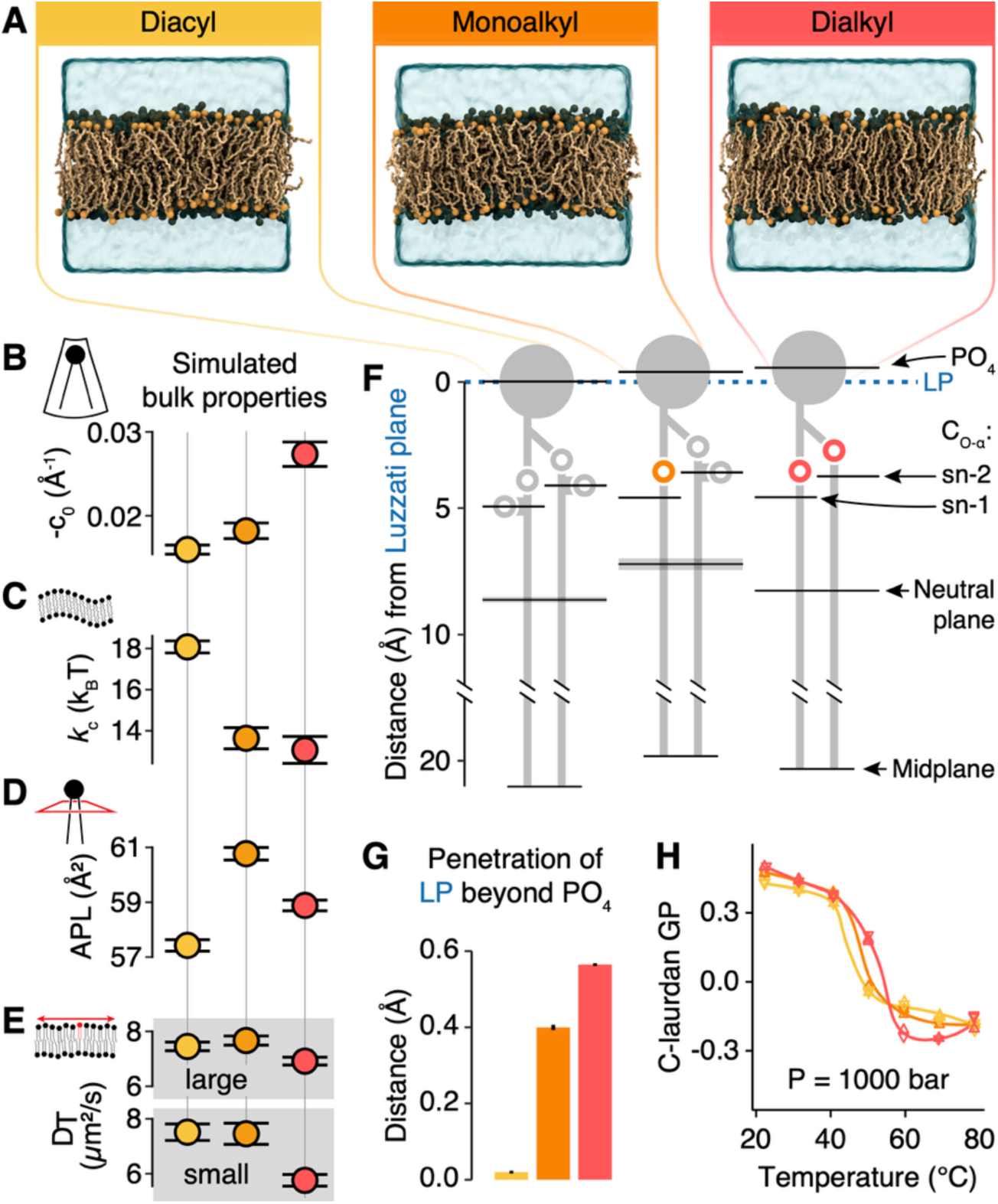
Atomistic simulations reveal a structural basis for the mechanical effects of alkyl linkages. (**A**) Bilayer simulations of the alkyl-POPE series at 35 °C and 1 bar. Bulk lipid properties derived from these simulations recovered trends similar to those in experimental data. (**B**) Alkyl linkages drive successive increases in −*c*_0_. (**C**) Introduction of the sn-1 alkyl linkage softens the bilayer. (**D**) Both alkyl species exhibit larger APL than diacyl, consistent with membrane thinning. (**E**) The dialkyl backbone decreases lateral diffusion in both large and small simulations. Monoalkyl has no effect. (**F**) Alkyl linkages remove steric bulk above the neutral plane and facilitate water penetration beyond the headgroup phosphate. Horizontal marks indicate transbilayer distances measured in simulations of key atoms and the neutral plane from the LP. C_O-α_ = carbon alpha to the glycerol oxygen. (G) Alkyl linkages enhance water penetration. (H) C-laurdan GP across the chain-melting transition at 1000 bar. Despite melting at a higher temperature, dialkyl PE exhibits lower GP in the L_α_ phase, implying greater water penetration. High-T increase in dialkyl GP indicates H_II_ formation. Error bars/bands in all panels are ±SEM.

Flexibility-fluidity decoupling was assessed in simulations via area per lipid (APL) and the translational diffusion coefficient *D_T_*. Alkyl backbones exhibited larger APL than the diacyl backbone (Fig. 4D, Table S5B), consistent with the thinner monolayers they formed in experiments and most simulations (Table S6B). However, the additive effect of ether linkages was not recovered. *D_T_*, which carries a finite-size dependence on simulation box dimension, was evaluated in both small and large (10×10 and 20×20 nm) systems; at both box sizes, dialkyl *D_T_* was lower than monoalkyl (Fig. 4E, Table S5B), which was indistinguishable from diacyl (P>0.35, Wald z-test). In sum, bilayer simulations corroborated the thinning and softening effects of ether linkages concomitant with negligible to negative impact on fluidity.

To compare aspects of lipid conformation along the transbilayer axis, we designated three specific atoms as “molecular fiducials” and computed their depth relative to solvent, the bilayer midplane, and the neutral plane of monolayer bending computed by TCB (Fig. 4F). We observed that the backbone linkages – specifically where an oxygen is removed from the carbonyl in the alkyl compared to acyl linkages – consistently lay on the headgroup side of the neutral plane. The removal of steric bulk 2.6-4.5 Å “above” this plane, where bending toward the headgroup involves lateral compression, might thus explain the additive curvature-enhancing effect of alkyl linkages observed in both experiments and simulations. Reduced linkage sterics corresponded with an increased penetration of water into the backbone region (past the PO_4_ group), as measured through position of the Luzzati Plane (LP) – the surface at which the water deficit outside balances the excess inside (Fig. 4G). This observation was consistent with C-laurdan fluorescence of liposomes composed from the same lipids, whose Generalized Polarization (GP) is a measure of water penetration into the membrane (70). GP was reduced in all alkyl-linked PEs compared to their diacyl analogs (Figs. 4H, S4B). This effect could not be explained by reduced ordering in the mono- and dialkyl systems, since the *T_m_* values determined by this method (Fig. S4B) were consistent with DSC (Fig. 3D) and were higher than that for diacyl POPE. Thus, both simulation and experiment indicate that alkyl ether linkages reduce lipid sterics above the neutral plane and admit solvent further into the backbone region of the bilayer.

### Plasmalogens promote the inverted hexagonal phase via two complementary mechanisms

Plasmenyl PE promotes H_II_ much more than dialkyl PE, despite displaying comparable intrinsic curvature (Fig. 2E). We therefore hypothesized that in addition to destabilizing the L_α_ phase via curvature elastic strain (CES), the plasmenyl linkage uniquely stabilizes H_II_ by relieving its interstitial packing frustration (IPF; Fig. 5A). To test the effects of backbone chemistry on these two free-energy contributions, we applied the model of Kirk et al. at excess hydration (71) to our experimentally derived phase behavior, −*c*_0_, and *k_c_* data. Under the Kirk framework, formation of H_II_ phase in the absence of an apolar filler requires lipid chains to stretch and splay non-uniformly to fill the interstitial spaces between adjacent tubules, incurring a largely entropic free-energy penalty we term IPF. This IPF cost can be reduced by decreasing the radii of the H_II_ tubes, but when the radius is smaller than the relaxed radius 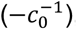, this incurs a CES penalty. These two contributions balance to determine the unit cell size of the H_II_ phase. CES also exists in the lamellar phase when the monolayers individually have nonzero intrinsic curvature. At the H_II_→L_α_ phase boundary, the free energy densities of the two phases are equal:

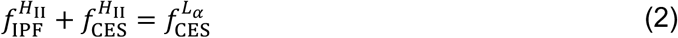

**Figure 5:**
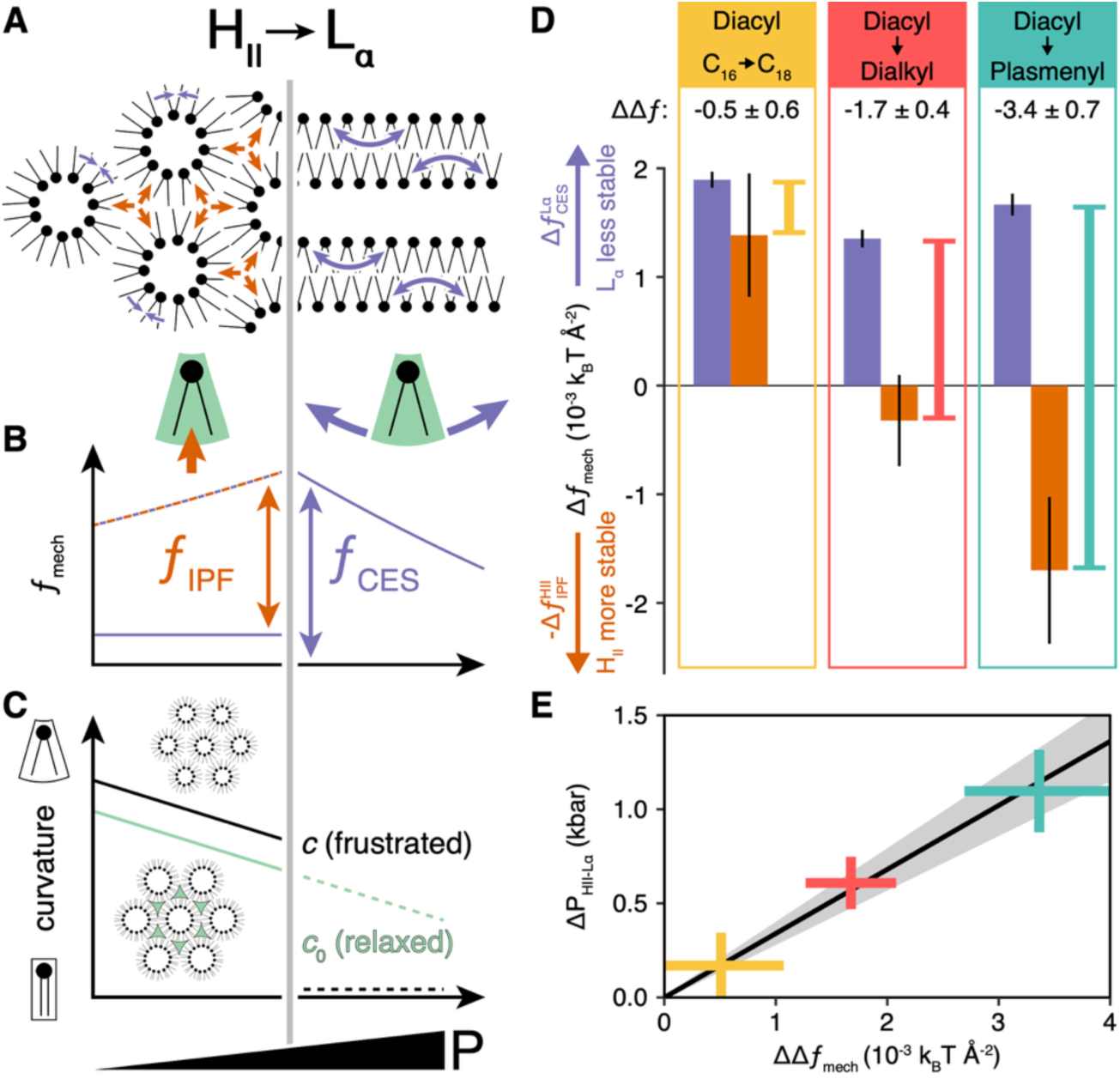
Mechanisms of H_II_ phase stabilization by dialkyl and plasmenyl PE. (**A**) Schematic of the barotropic H_II_→L_α_ transition illustrating areal free energy density (f) due to interstitial packing frustration (IPF, orange) and curvature elastic strain (CES, purple). Under this model, −*f*_IPF_is the major determinant of H_II_ stability and −*f*_CES_ is the major determinant of L_α_ stability. (**B**) Schematic of mechanical free energy components around the H_II_-L_α_ transition. At the transition, (*f*_IPF_ + *f*_CES_, orange/purple dashed line) of H_II_ is equal to *f*_CES_ of L_α_ (purple line to the right of transition). Measurement of *f*_CES_ in both H_II_ and L_α_ therefore enables isolation of *f*_IPF_. (**C**) Schematic of observable curvature trends used to calculate *f*_CES_. Light green represents apolar hydrocarbon used to relax the H_II_ lattice. Intrinsic curvature is extrapolated into the L_α_ regime; mean *c* is assumed to be zero in bilayers. (**D**) Effects of lipid structural substitutions on energetic contributions to inversion. Purple bars indicate CES destabilization of L_α_ and orange bars IPF stabilization of H_II_, with ±SEM error bars. Yellow, red, and teal lines indicate *ΔΔf*_mech_ of inversion associated with each substitution; numerical values are reported at the top of each panel. (**E**) Differences in pressure required to drive pairs of lipids from H_II_→L_α_ at 75 °C (*P_H_*_II→*L*_*_α_*) track with *ΔΔf*_mech_ in each pair of lipids.

All energies here are referred to the pivotal plane, whose area per lipid is by definition unchanged by bending; taking this area to be the same in both phases, it cancels and the coexistence condition reduces to an equality of areal free-energy densities, equivalent to the per-lipid comparison of Kirk et al. (58).

As *f*_CES_ depends only on intrinsic curvature and bending modulus (Eq. 1), our measurements of *c*_0_(*P*) and *k_c_* allowed the CES contributions in both phases to be constrained. The IPF contribution to H_II_ stability 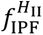 was then obtained by difference at the phase transition (Fig. 5B) using *c* and *c*_0_ evaluated at that transition (Fig. 5C). For every pair of lipids compared, the difference in the curvature elastic strain energy density of the lamellar phase 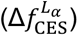 was evaluated at a common reference pressure (Methods, Eqs S35-S38; Fig. S6C). This analysis revealed distinct energetic effects of structural substitutions on these two parameters (Fig. 5DE). Because it is recovered by difference, the IPF term encompasses every contribution to inverted-phase stability not captured by CES. Strictly, it represents an upper bound on the energetic role of chain packing. In our use of the Kirk model, we chose to omit contributions from electrostatics and interlamellar interactions because (i) our systems were free of charged lipids and (ii) we estimated the free energy densities of interlamellar interactions to be roughly an order of magnitude smaller than the backbone-dependent differences resolved here (Fig. S7; SI Appendix, Role of interlamellar interactions).

For comparison, we also considered a non-backbone substitution. Extending the *sn-*1 chain of diacyl PE from 16:0 to 18:0 produced a large increase in 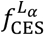 but it also raised 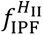 by a comparable amount, so that the difference between these contributions was insignificant (Fig 5D, first panel). Substituting the dialkyl for the diacyl backbone also augmented 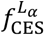, but with no IPF penalty destabilizing H_II_, to give *ΔΔf*_mech_ = -1.67±0.41 x10^-3^ *k_B_T*/Å^2^, favoring H formation. The plasmenyl backbone exhibited similar destabilization of L_α_ and additionally stabilized H_II_, resulting in *ΔΔf*_mech_ of a large magnitude favoring H_II_ (-3.37±0.67 x10^-3^ *k_B_ T*/Å^2^). Taken together, values calculated for *ΔΔf*_mech_ closely track the observed shifts in inversion pressure (Fig. 5D, E). These results indicate that ether lipids can stabilize non-lamellar topologies through two separable mechanisms: (1) negative intrinsic curvature, which is shared by alkyl and plasmenyl linkages, and (2) relief of interstitial packing frustration, which is unique to plasmalogens. These two mechanisms working in concert account for the exceptional ability of plasmenyl PE to promote inverted phase formation.

## Discussion

The results presented here demonstrate phospholipid backbone-linkage chemistry as a modular control element for membrane mechanics that allows intrinsic curvature, bending stiffness, and non-lamellar phase propensity to be tuned independently of membrane fluidity and hydrocarbon chain order. The two types of ether linkages – alkyl and plasmenyl – confer distinct combinations of membrane-mechanical properties (Fig. 6), each of which is likely important in certain biological contexts. The measured effects of linkage type on the properties of the comparative PE lipids is showing in Table 1.

**Figure 6:**
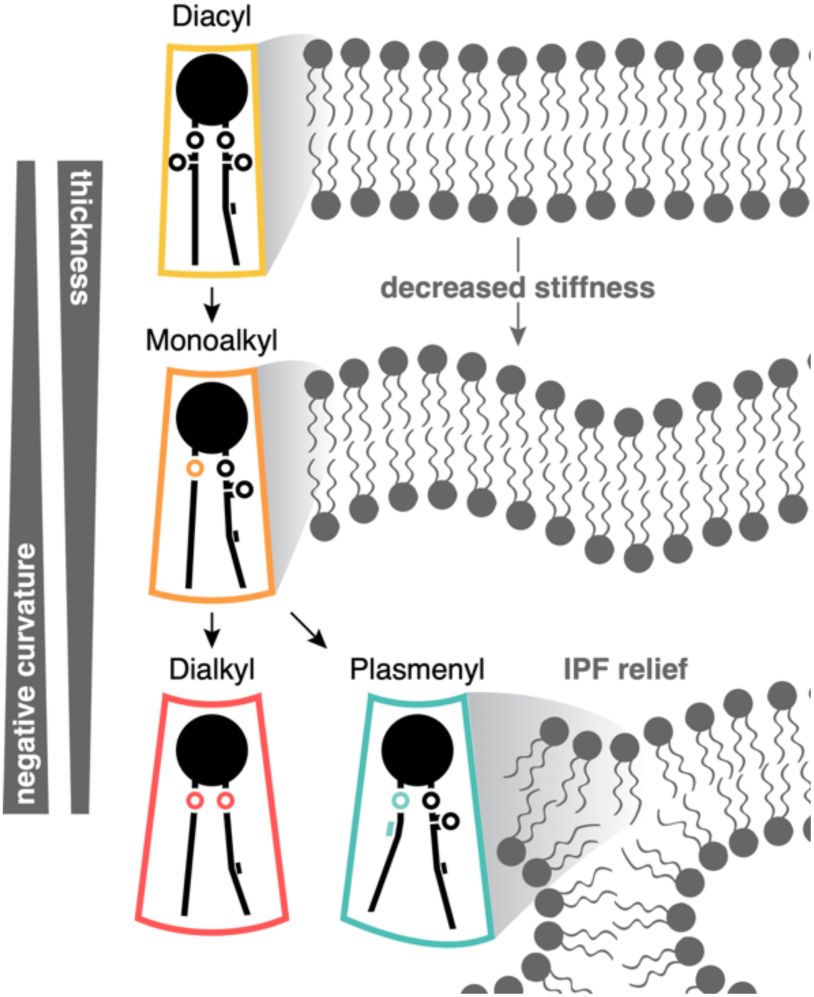
The membrane-mechanical code of backbone linkage chemistry. The substitution of alkyl for acyl linkages increases negative intrinsic curvature and decreases mono/bilayer thickness and stiffness. Addition of the plasmenyl C=C bond to the sn-1 alkyl chain augments these effects and relieves interstitial packing frustration to stabilize the H_II_ phase.

**Table 1.** Backbone-linkage effects on membrane mechanical parameters and melting temperature. *Δf* values are changes vs. the corresponding diacyl reference at the barotropic H_II_→L_α_ transition (75 °C). (N/D), not determined.

| Lipid | $-c_0 \pm \text{SEM}$<br>(Å<sup>-1</sup>) | | $k_c \pm \text{SEM}$<br>($k_B T$) | $\Delta f_{\text{CES}}^{L_{\alpha}} \pm \text{SEM}$<br>(10<sup>-3</sup><br>$k_B T/\text{Å}^2$) | $\Delta f_{\text{IPF}}^{H_{\text{II}}} \pm \text{SEM}$<br>(10<sup>-3</sup><br>$k_B T/\text{Å}^2$) | $T_m \pm \text{FWHM}$<br>(°C) |
| --- | --- | --- | --- | --- | --- | --- |
|  | 35 °C | 75 °C |  |  |  |  |
| diacyl POPE | 0.02213<br>± 6.7×10<sup>-4</sup> | 0.035028<br>± 6.2×10<sup>-5</sup> | 10.324<br>± 0.084 | (reference) |  | 25.66<br>± 1.3 |
| monoalkyl POPE | 0.02392<br>± 3.2×10<sup>-4</sup> | 0.03791<br>± 7.6×10<sup>-5</sup> | 8.416<br>± 0.072 | (N/D; forms Q<sub>II</sub>) |  | 26.89<br>± 0.92 |
| dialkyl POPE | 0.03073<br>± 4.6×10<sup>-4</sup> | 0.040296<br>± 6.2×10<sup>-5</sup> | (N/D) | 1.35<br>± 0.08 | -0.32<br>± 0.42 | 27.67<br>± 1.2 |
| diacyl SOPE | 0.02308<br>± 3.1×10<sup>-4</sup> | 0.035424<br>± 8×10<sup>-5</sup> |  | (reference) |  | 30.89<br>± 1.1 |
| plasmenyl SOPE | 0.02645<br>± 8.2×10<sup>-4</sup> | 0.040765<br>± 7.6×10<sup>-5</sup> |  | 1.67<br>± 0.10 | -1.70<br>± 0.68 | 27.73<br>± 0.73 |

The replacement of acyl with alkyl linkages increases negative intrinsic curvature and thus the CES energy stored in flat bilayers. This destabilization of L_α_ alone drives a substantial enhancement of H_II_ formation and hemifusion. The plasmenyl linkage introduces the additional, distinct effect of relieving IPF energy stored in H_II_, giving plasmalogens their exceptional H_II_ propensity and fusogenicity. This is consistent with prior proposals that plasmalogens could promote inversion by stabilizing H_II_ (65) through extension of the sn-2 chain that was observed by NMR (71). In our decomposition, this sn-2 extension appears as reduced IPF: the vinyl-ether linkage’s near-planar interfacial geometry plausibly allows the sn-2 chain to fill interstitial volume in the inverted lattice at lower energetic cost. This is analogous to the role hydrocarbons like TS impart on H_II_ stability. We amend this model by finding that the plasmenyl – as well as the alkyl – linkage does in fact destabilize L_α_ through increased negative intrinsic curvature. The simultaneous L_α_ destabilization and H_II_ stabilization is a property of plasmalogens not found in other ether lipids. Under our experimental conditions, the 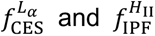 contributions to H_II_ propensity by plasmenyl PE were similar in magnitude.

Through systematic comparison, we discovered that backbone linkage chemistry also modulates a pair of membrane properties – CES and bending stiffness – that are otherwise difficult to tune independently. Substitution of ester with ether linkages increases negative monolayer spontaneous curvature while softening the membrane, yet has comparatively modest effects on hydrocarbon packing; any residual changes can be readily offset by adjusting acyl chain unsaturation. Atomistic simulations corroborate this decoupling: even as the bending modulus fell, simulated lateral diffusion did not increase and area per lipid tracked the measured thinning (Fig. 4D,E). In this sense, ether linkages provide a compact, orthogonal means of regulating membrane topology change. Other common strategies are more strongly coupled. Headgroup substitution or removal can shift curvature but also substantially alters hydration, ion interactions, and lateral organization (72). Modifying chain unsaturation or length likewise perturbs curvature and elasticity, but does so in tandem with fluidity and hydrocarbon packing.

Pairing the plasmenyl linkage with the PE headgroup (as opposed to distributing it between PE and PC) yields a species with a distinctly strong negative *c*_0_ and so facilitates reshaping through its dynamic enrichment at highly curved regions of transition structures – an effect that scales as the square of *c*_0_ (61). Beyond its impact on curvature elastic strain, the plasmenyl linkage further stabilizes non-lamellar topology through relief of interstitial packing frustration, an effect that may also be achieved by chain-length mismatch but again at the cost of pronounced packing perturbations (73). Cholesterol and related sterols can modulate elastic properties, but can induce lateral phase separation (74). Finally, the distinct phase behavior of monoalkyl lipids – intermediate in −*c*_0_ and *k_c_* yet favoring the Q_II_ phase – indicates that alkyl and plasmenyl linkages are not functionally interchangeable, likely reflecting differences in Gaussian curvature contributions that are mitigated by the plasmenyl C=C bond. These varying features – intrinsic and Gaussian curvatures for dialkyl and monoalkyl ether linkages, as well as relief of interstitial packing frustration for plasmenyl linkages – likely explain their similar effects on fusion of model membranes, a complex process dependent on multiple mechanical properties (Extended Discussion).

Although our analyses focused on bulk behavior, they also suggest molecular mechanisms by which subtle linkage chemistry changes control membrane properties. We propose a simple steric mechanism for the enhancement of −*c*_0_, and thereby *Δf*_CES_, by alkyl linkages. The removal of ester carbonyl groups entails a reduction in area per lipid above the neutral plane of bending (75), which appears to directly drive a more tapered lipid shape as acyl→alkyl substitutions are made. We considered the alternative model that water mediates the change in cross-sectional area through hydrogen bonding with ester carbonyls, but neither atomistic simulations (Fig. 4) nor fluorescence spectroscopy experiments (Fig. S4) recovered hydration patterns consistent with this mechanism. As acyl linkages were replaced with alkyl, the Luzzati plane in L_α_ simulations consistently moved deeper into the membrane relative to the phosphate group, and C-laurdan GP decreased, indicating greater water penetration. The latter result agreed with previous laurdan analysis of alkyl ether lipids (70). Increased water penetration through the headgroup layer increases lipid spacing, thins the membrane, and lowers *k_c_*.

The repeated evolution and horizontal transfer of alkyl and plasmenyl ether lipid biosynthesis suggests a broad functional basis for incorporation of these lipids into cell membranes beyond that of chemical stability or roles in oxidative signaling and stress. More recently, we implicated plasmalogens in the adaptive maintenance of curvature elastic strain in high-pressure deep-sea environments (46), but comparably large fractions of plasmalogen are found in specific mammalian tissues. Mammalian cells increase ether lipid content in response to pressure-induced reduction of −*c*_0_, suggesting that their metabolism regulates membrane CES (76). Other work has recently highlighted the distinct trafficking routes that ether phospholipids take compared to their diacyl counterparts (76). Our results show that ether linkages impart specific biophysical properties to phospholipids: (1) destabilization and (2) softening of bilayers as well as (3) stabilization of the fusion/fission-relevant H_II_ phase upon desaturation to a plasmenyl linkage. By decoupling stiffness, intrinsic curvature, and IPF from chain packing, backbone linkage chemistry could enable cells to tune membrane topology changes and CES in response to perturbation in ways that headgroup and chain modifications cannot. The unique combination of membrane-mechanical properties conferred by ether linkages is likely relevant in explaining their distribution across bacteria and eukaryotes, as well as their structural context within lipidomes.

## Materials and Methods

The Supplementary Information contains detailed protocols for all experiments and analyses. Experimental methods include vesicle fusion assays and validation, collection and analysis of SAXS data, bending moduli measurements by SAXS and micropipette aspiration/optical tweezers, DSC and laurdan GP measurements. Computational analyses covered include phylogeny representation, molecular dynamics simulations, and extraction of structural, dynamic, and mechanical parameters from simulations.

## Acknowledgements

Sol Gruner and Maria Fedorova contributed key discussions. Beamline scientists Estella Yee, Qingqiu Huang, and Richard Gillilan provided technical assistance. Research was supported by funding from the Office of Naval Research (N00014-23-1-2543 to I.B.), the National Institutes of Health (5T32EB009380-14 to D.M., GM142960 to I.B., 1ZIAHD008955 to A.J.S.), the NASA Postdoctoral Program Fellowship program (0017-NPP-MAR22-A-Astrobio to J.R.W.), the Gordon and Betty Moore Foundation (9208 to P.R.G.), and the National Science Foundation (MCB-2316457 to E.L., MCB-2046303 and MCB-2316458 to I.B., and DEB-2321651 to P.R.G.). SAXS measurements were performed at the Center for High-Energy X-ray Sciences (CHEXS), which is supported by NSF award DMR-2342336, and the MacCHESS resource is supported by NIH award 1-P30-GM124166.

## Notes

https://github.com/octopode/etherlipids-2026

### Competing Interest Statement

The authors have declared no competing interest.

### Summary of Updates

This version of the manuscript has been revised based on peer feedback; additional computational and experimental results are presented in the main text and Supporting Information. One coauthor has been added.

