## Supporting Information for "Ether linkages in phospholipids provide modular control of membrane mechanics"

**This PDF file includes:**

Methods
Extended Discussion
Figures S1 to S7
Tables S1 to S6
SI References

**Supporting Text**

**Methods**

Phylogenetic distribution and structural context of ether backbone linkages by meta-analysis

***Ether lipid biosynthesis across the tree of life***

Genomic annotations for the four known committed-step enzymes of ether lipid biosynthesis were collected from the literature. These included the anaerobic-pathway enzymes Ger (1) and PlsR (2, 3), as well as the aerobic-pathway enzymes AGPS (1) and CarF (4, 5). Annotations were mapped to bacteria in the Global Taxonomy DataBase (GTDB) using its Taxonomic Name Resolution Service (6) and low-confidence matches were manually resolved. The GTDB tree was collapsed to class level and each class was scored for presence or absence of each of the four enzymes within the clade. Evidence of gain or loss of plasmalogen biosynthesis in stem-group eukaryotes was collected from the literature (5, 7) and mapped manually onto the Open Tree of Life (OpenTOL) topology (8) using the induced subtree function of the OpenTOL REST API. Phylogenies were plotted in R using ggtree (9); the eukaryote tree was manipulated manually to fit the figure layout.

***Headgroup structural context of backbone types in metazoans***

Phospholipid backbone compositions under the PE and PC headgroups were collected from the literature. We used only source publications that unambiguously resolved monoalkyl and plasmenyl linkages using sequential hydrolysis, liquid chromatography, or $MS^2$ fragmentation. Mass abundances of backbone types under each headgroup were normalized to a sum of 1.

Assay of calcium-catalyzed vesicle hemifusion kinetics

***Vesicle hemifusion lipid mixing FRET assay***

A FRET-based lipid mixing assay was performed by fluorescence dequenching of fusing liposomes with equimolar POPC, POPS, and varying PE species. As a negative control, a condition with only equimolar POPS and POPC was tested. In this assay, two populations of liposomes are mixed; one with both 2 mol% FRET acceptor (Rh-DPPE, Avanti Research) and FRET donor (NBD-DPPE or NBD-PE, Avanti Research) and the other with no fluorophores. A lipid film for equimolar lipid concentrations (200μM final) was prepared from lipid stocks in chloroform and evaporation of the organic solvent under $N_2$ for 5 minutes and then held in a vacuum chamber for an hour or more. Large unilamellar vesicles were prepared by rehydration in Buffer A (100mM NaCl, 5mM Na HEPES, 0.1mM EDTA, pH 7.4) with 30 minutes of tumbling, 4 freeze-thaw cycles, and extrusion through a 100nm membrane 21 times. Both the FRET pair containing and unlabeled vesicle groups were mixed in equal volumes and loaded in a quartz cuvette. Following the addition of 10mM $CaCl_2$ to induce fusion, the emission of NBD-PE dequenching (ex/em 463/536 nm) was measured using a fluorescence spectrophotometer (Cary Eclipse) in Kinetics mode for 5 minutes. Fluorescence of a fully dequenched control with 1 mol% NBD-DOPE and Rh-DOPE was measured for determination of total % fused vesicles.

***Measurement of vesicle lamellarity by dithionite quenching of outer leaflet NBD-PE***

Pre- and immediately post-fusion vesicle lamellarity was determined by bleaching outer leaflet NBD-PE with sodium dithionite. As previously described, two populations of vesicles were mixed (200 μM final) containing either equimolar diacyl POPE:POPC:POPS or monoalkyl POPE:POPC:POPS with or without 2% NBD-PE. The assay was performed before or after the addition of 10 mM $CaCl_2$, and vesicles were excited at 460 nm and emission was detected at 538 nm. At 30 s, 10 mM sodium dithionite (1 M stock dissolved in unbuffered 0.5 M Tris immediately before use) was added. At 3.5 min, 0.1% TritonX-100 was added to solubilize the vesicles and allow dithionite to quench the remaining NBD-PE. % NBD Emission Quenched is defined as (1-

(F-F0)/(Fmax-F0)) * 100, where Fmax is the original fluorescence intensity, F is the emission after dithionite addition, and F0 is signal remaining after vesicle solubilization with 0.1% Triton X-100. Experiments were conducted at 37 °C.

***Measurement of vesicle size by dynamic light scattering***
The hydrodynamic radius (nm) of vesicles was measured by DLS using a DynaPro 99-E-50 instrument with a temperature-controlled microsampler (Protein Solutions) with Dynamics software (Protein Solutions). Measurements were performed on vesicles following extrusion before fusion was initiated by 10 mM $CaCl_2$, 5 minutes post fusion, and 5 hours post fusion Cuvettes were cleaned with 70% ethanol, water, and drying between each measurement, and triplicate measurements were performed for each condition. Each condition measurement consisted of an average of 13 scans.

Construction of lipid barotropic phase diagrams and determination of $H_{II}$ dimensions by SAXS

***Sample preparation and data acquisition***
Lipid dispersions were prepared in excess pure water as previously described (10). For DOPE-hosted preparations, lipids (Avanti Polar Lipids) were purchased in $CHCl_3$ solutions of known concentration and mixed at the appropriate molar ratio using Wiretrol™ glass micropipettes (Drummond). For partially or fully relaxed preparations, 6 or 12% w/w 9(Z)-tricosene (Sigma-Aldrich) was added similarly. Samples were mounted in the high-pressure SAXS cell (11) using laser-cut PMMA sample holders fitted with either 7 µm thick polyimide windows (Chemplex) or a closed-end glass capillary (Kimble Chase). SAXS data were collected as previously described (10). Three 1 s exposures were taken every 100 or 200 bar with 60 s equilibration per 100 bar.

***Phase identification and fitting of repeat spacing***
Lipid phases were identified manually by specifying a scattering vector $q$ for the first-order Bragg peak of each phase signature present in each SAXS profile. Higher-order peaks were assigned to phases using characteristic spacings for the lattice geometry of each phase with a $q$-tolerance window. Where multiple peaks were available for a single phase, repeat spacing was estimated by least-squares fitting to expected peak positions and uncertainties were propagated from peak position errors.

Characteristic $q$ spacings used for each phase were:
$L_\alpha$ / $L_\beta$: 1, 2, 3, 4, 5, 6 …
$Q_{II}$ (Pn3m): 1, $\sqrt{1.5}$, $\sqrt{2}$, $\sqrt{3}$, 2, $\sqrt{4.5}$ …
$H_{II}$: 1, $\sqrt{3}$, 2, $\sqrt{7}$, 3, $\sqrt{12}$ …

***Determination of barotropic phase transitions***
Barotropic phase transitions were determined from SAXS pressure series by first estimating phase fractions from first-order peak intensities. For each series, first-order peak intensities for each phase were averaged across replicate exposures at each pressure stop, then normalized to that phase's maximal intensity across the experiment. These values were then renormalized across phases at each pressure so that total phase fraction summed to 1. To fit individual transitions, cumulative phase fractions were constructed in pressure order after excluding $H_{II}$, such that each non-$H_{II}$ phase reported the fraction of sample converted out of $H_{II}$ into that phase or any phase above it; this accommodated the analysis of 3 coexisting phases as observed in monoalkyl PE. Curves were trimmed to retain a single zero-valued point at the low-pressure edge and to remove non-monotonic points within a sweep. Transition pressures were then obtained by fitting these stacked fractions versus pressure with three-parameter binomial logistic models. The two pressure directions were weighted to contribute equally when sweep lengths differed and thus

balance hysteresis. The inflection point of each logistic curve was taken as the transition pressure and its standard error was obtained from the fitted model.

***Determination of monolayer curvature $c$ and effective hydrocarbon thickness $h_{\mathrm{eff}}$***

Multiple methods exist for determining $c$ from $H_{II}$ SAXS data, though they yield values defined at different planes within the monolayer. Electron-density models fitted globally to SAXS profiles (12, 13) resolve curvature at the neutral plane, the depth at which bending and stretching are energetically decoupled (14). Geometric analyses, using $H_{II}$ lattice dimensions and partial molar volumes (14, 15), determine curvature at the pivotal plane, the depth at which the lipid cross-sectional area is unchanged by bending. Though the neutral plane is considered the more mechanically rigorous reference plane, both planes lie near the glycerol backbone and their depths tend to differ by ≤10% (14). In PE monolayers, the pivotal plane lies slightly further from the headgroup than the neutral plane (14–17), thus returning slightly smaller $-c_0$. As we were not able to non-degenerately constrain all the necessary parameters of the $H_{II}$ global scattering model, we used the geometric method described below, which recovers curvature at the pivotal plane as well as mechanical effective hydrocarbon thickness $h_{\mathrm{eff}}$, defined as the distance from the pivotal plane to the chain termini.

The PC and SOPE series studied here do not form $H_{II}$ at 35 °C, motivating our uniform use of the host-guest method to measure $c_0$ at this temperature. This entailed doping each "guest" lipid to be measured into a diacyl-DOPE "host" matrix (PE at 20 mol %, PC at 10 mol %) and assuming curvature averages linearly with guest fraction (18). Guest fractions were kept ≤20 mol % to avoid $H_{II}$ destabilization and potential demixing artifacts. Unhosted lipid samples were measured uniformly at 75 °C to make the barotropic $H_{II}$→$L_{\alpha l}$ transition accessible and thus enable the decomposition of energetic contributions to this phase transition (see below.)

For each sample, the first-order $H_{II}$ peak position $q$ was converted to layer spacing $d$:

$$d = \frac{2\pi}{q} \tag{S1}$$

And lattice parameter $a$:

$$a = \frac{2d}{\sqrt{3}} \tag{S2}$$

Rather than estimating monolayer geometry from a Luzzati-plane construction, we solved for the pivotal plane radius using literature-derived hydrocarbon volumes together with a geometric saturation model for tricosene incorporation. Lipid hydrocarbon volume was represented by the DOPE-referenced scheme,

$$V_{\mathrm{hc},j}^{\mathrm{ref}} = V_{\mathrm{hc,DOPE}}^{\mathrm{ref}} + \Delta n_{\mathrm{CH_2},j}(2V_{\mathrm{CH_2}}) + \Delta n_{\mathrm{db},j}\Delta V_{\mathrm{db}} + \Delta n_{\mathrm{ether},j}\Delta V_{\mathrm{ether}}, \tag{S3}$$

with

$$\Delta V_{\mathrm{db}} = V_{\mathrm{CH=CH}} - 2V_{\mathrm{CH_2}}, \tag{S4}$$

and mixed-sample values were taken as host-guest mole-fraction averages. Pivotal-plane areas were treated analogously, with PE reference values $A_{p,\mathrm{PE}}^{0}$ and $A_{p,\mathrm{PE}}^{\mathrm{sat}}$, and a fixed PC offset $\Delta A_{p,\mathrm{PC}}$ added to both. Central values were $A_{p,\mathrm{PE}}^{0} = 64.2$ Å$^2$, $A_{p,\mathrm{PE}}^{\mathrm{sat}} = 73.6$ Å$^2$, and $\Delta A_{p,\mathrm{PC}} = 2.0$ Å$^2$ (15, 19). The hydrocarbon reference volume for DOPE, the methylene-pair increment, the olefinic increment, and the ether correction were taken from the lipid-volume literature (20–23). Temperature and hydrostatic pressure were incorporated through separate volume and area scaling factors,

$$S_V = 1 + \alpha_V(T - T_{\mathrm{ref}}) - \kappa_V(P - P_{\mathrm{ref}}) \tag{S5}$$

and

$$S_A = 1 + \alpha_A(T - T_{\mathrm{ref}}) - \kappa_A(P - P_{\mathrm{ref}}), \tag{S6}$$

with $T_{\mathrm{ref}} = 20\ ^\circ\mathrm{C}$, $P_{\mathrm{ref}} = 1\ \mathrm{bar}$, $\alpha_V = 1.0 \times 10^{-3}\ K^{-1}$, $\kappa_V = 4.5 \times 10^{-5}\ \mathrm{bar}^{-1}$, $\alpha_A = 2.5 \times 10^{-3}\ K^{-1}$, and $\kappa_A = R_{\mathrm{aniso}}\kappa_V$, using an area-volume compressibility ratio $R_{\mathrm{aniso}} = 3.0$ in the central analysis (20). Thus $V_{\mathrm{hc}}^{\mathrm{lipid}} = V_{\mathrm{hc}}^{\mathrm{mix,ref}} S_V$, while $A_p^0$ and $A_p^{\mathrm{sat}}$ were multiplied by $S_A$.

The added TS volume per phospholipid was computed from the input mass fraction $w_{\mathrm{TS}}$ as

$$V_{\mathrm{TS}}^{\mathrm{add}} = \frac{w_{\mathrm{TS}}}{1 - w_{\mathrm{TS}}} \cdot \frac{M_{\mathrm{mix}}}{M_{\mathrm{TS}}} \cdot \bar{v}_{\mathrm{TS}} \cdot S_V, \tag{S7}$$

with $M_{\mathrm{TS}} = 322.6\ \mathrm{g\ mol}^{-1}$ and $\bar{v}_{\mathrm{TS}} = 670\ \mathring{A}^3$ per molecule.

The pivotal-plane radius $r$ was then obtained by solving the $H_{\mathrm{II}}$ unit-cell volume balance (24, 25). Defining

$$A_{\mathrm{cell}} = \frac{\sqrt{3}}{2} a^2, \tag{S8}$$

$$A_{\mathrm{ann}}(r) = \pi\left[\left(\frac{a}{2}\right)^2 - r^2\right], \tag{S9}$$

and

$$A_{\mathrm{int}} = a^2\left(\frac{\sqrt{3}}{2} - \frac{\pi}{4}\right), \tag{S10}$$

the maximum incorporable TS volume per phospholipid was

$$V_{\mathrm{TS}}^{\mathrm{max}}(r) = V_{\mathrm{hc}}^{\mathrm{lipid}} \frac{A_{\mathrm{int}}}{A_{\mathrm{ann}}(r)}, \tag{S11}$$

the incorporated TS volume was

$$V_{\mathrm{TS}}^{\mathrm{inc}}(r) = \min\left(V_{\mathrm{TS}}^{\mathrm{add}}, V_{\mathrm{TS}}^{\mathrm{max}}(r)\right), \tag{S12}$$

and the associated saturation fraction (capped at 1 per Eq. S12) was

$$\theta(r) = \frac{V_{\mathrm{TS}}^{\mathrm{inc}}(r)}{V_{\mathrm{TS}}^{\mathrm{max}}(r)}. \tag{S13}$$

The effective pivotal-plane area was then interpolated as

$$A_p(r) = A_p^0 + \theta(r)\left(A_p^{\mathrm{sat}} - A_p^0\right), \tag{S14}$$

and $r$ was defined implicitly by

$$\frac{A_{\mathrm{cell}} - \pi r^2}{2\pi r} = \frac{V_{\mathrm{hc}}^{\mathrm{lipid}} + V_{\mathrm{TS}}^{\mathrm{inc}}(r)}{A_p(r)}. \quad \text{(S15)}$$

This equation was solved numerically for each $H_{II}$ phase SAXS exposure. Monolayer curvature and effective hydrocarbon thickness were computed on the same pivotal surface as

$$c = -\frac{1}{r} \quad \text{(S16)}$$

and

$$h_{\mathrm{eff}} = -r + \sqrt{r^2 + \frac{2V_{\mathrm{hc}}^{\mathrm{lipid}} r}{A_p(r)}}. \quad \text{(S17)}$$

In relaxed samples containing sufficient TS and no applied osmotic stress, $c$ represents the intrinsic curvature $c_0$.

Because monolayer curvatures were used to separate energetic components of phase stability (see below) and hydrostatic pressure was used to drive phase transitions in SAXS experiments, curvatures were measured across P sweeps at a constant T.

The $c$ vs. $P$ relationships observed were generally linear with similar slopes, so a linear model was fitted jointly to the P-series of all unhosted PE systems. The model chosen used the formula

$$c = a_j\{lipid\colon ts = j\} + \beta P \quad \text{(S18)}$$

where $ts$ is the tricosene (TS) fraction, $\alpha_j$ is the intercept for each lipid-TS combination, and $\beta$ is a global slope with pressure. This parameterization was favored by AIC over models treating TS as a uniform linear or categorical effect because the influence of TS differed among lipids; a fully interacted lipid×TS model gave equivalent information content but less compact parametrization. The selected structure was fitted by robust linear regression using statsmodels RLM, using the Huber T M-estimator and H1 covariance due to heavy tails in the OLS residual distribution. A robust linear model was also used to fit $h_{\mathrm{eff}}$, with the formula

$$\log(h_{\mathrm{eff}}) = \alpha c(\mathrm{lipid}, \mathrm{ts}, P) + \beta P + \gamma P^2 \quad \text{(S19)}$$

**Sensitivity analysis**
A tiered deterministic sensitivity analysis was used to account for uncertainty in the literature-derived parameters. Tier 1 varied each uncertain parameter individually to predefined low and high values and recorded the resulting changes in $c$ and $h_{\mathrm{eff}}$. The screened parameters were $A_{p,\mathrm{PE}}^{0}$, $A_{p,\mathrm{PE}}^{\mathrm{sat}}$, $\Delta A_{p,\mathrm{PC}}$, $\alpha_V$, $\kappa_V$, $\alpha_A$, $R_{\mathrm{aniso}}$, TS molar volume, $V_{\mathrm{hc,DOPE}}^{\mathrm{ref}}$, $V_{\mathrm{CH_2}}$, $V_{\mathrm{CH=CH}}$, and $\Delta V_{\mathrm{ether}}$. Tier 2 then selected the top four Tier 1 drivers, ranked by maximal absolute percent effect in hosted 35 °C samples, and evaluated all pairwise low/high combinations to detect interactions and changes in TS-saturation regime. These analyses are reported in Tables S2 and S3 as deterministic response bands and influence rankings.

<u>Measurement of PE monolayer bending moduli by SAXS</u>

***Sample preparation and data acquisition***

Solutions of 15 and 60% w/v PEG 20000 (Sigma-Aldrich) were prepared in pure water (Low-TOC HPLC grade, MilliporeSigma). Osmolality $m_{\mathrm{osm}}$ of these solutions was measured at 30.1°C using a vapor pressure osmometer (Wescor Vapro 5600); osmotic pressure $\pi$ at 75 °C was obtained from the van 't Hoff equation:

$$\pi_{348.15K} = R(348.15K) \cdot m_{\mathrm{osm}348.15K}. \quad \text{(S20)}$$

Lipids were prepared with 12% w/w 9(Z)-tricosene (Sigma-Aldrich) and dispersed in excess pure water as previously described (10). Three samples of each lipid were prepared in closed-end glass capillaries (Kimble Chase), which were cut to 15 mm length, then heated to 75 °C in a water bath. The water above the hydrated lipid paste was exchanged 3 times with preheated water (control) or PEG solution using a syringe and needle. Samples were capped with high-vacuum grease (Corning) to prevent evaporation, then equilibrated 2 h in a 75 °C bath. Each equilibrated sample was mounted in the high-pressure sample holder and SAXS data collected as previously described (10). Three 1 s exposures were taken every 200 bar with 120 s equilibration per pressure step. When a phase change became apparent (near 1000 bar), the pressure sweep was stopped and the sample exchanged. Only pressure up-sweeps were collected because the monolayer topology change associated with an $H_{II} \rightarrow L_{\alpha}$ or $H_{II} \rightarrow Q_{II}$ phase transition irreversibly compromises the osmotic gradient.

***Derivation of monolayer bending rigidity from $H_{II}$ repeat spacing***

Hexagonal repeat spacing was obtained from the first-order $H_{II}$ Bragg peak and pivotal-plane radius $r$ was calculated as described above. Monolayer bending rigidity was then estimated from PEG-induced deformation of the $H_{II}$ lattice. For each stressed sample, the reference radius $r_0$ was defined as the mean pivotal-plane radius measured for the same lipid at the same hydrostatic pressure in the absence of PEG. To facilitate regression analysis, we calculated the theoretically linear deformation parameter

$$g = \frac{(1/r - 1/r_0)}{r^2}, \quad \text{(S21)}$$

with units of $\mathrm{\mathring{A}}^{-3}$ (26). All $(\pi, g, P)$ data with $\pi > 0$ were fit by weighted least squares, with weights set to $\mathrm{SEM}(\pi)^{-2}$. We compared several candidate models for $\pi(g, P)$, including one with a shared hydrostatic pressure effect and a fully interacted model with [PEG] as a random factor. The model best supported by AICc allowed PEG-specific diacyl baselines and P-effects, while constraining the monoalkyl effect to a shared fractional perturbation varying linearly with P. This parameterization also offered a single interpretable softening factor for the monoalkyl backbone. In this model,

$$k_c^{\mathrm{diacyl}}(P, 15) = \alpha_{15} + \beta_{15} P, \quad \text{(S22)}$$

$$k_c^{\mathrm{diacyl}}(P, 60) = \alpha_{60} + \beta_{60} P, \quad \text{(S23)}$$

where subscripts represent [PEG] (% w/v).

Monoalkyl-to-diacyl modulus ratio was written as

$$q(P) = q_0 + q_1 P. \quad \text{(S24)}$$

Thus,

$$k_c^{\mathrm{monoalkyl}}(P, 15) = q(P) k_c^{\mathrm{diacyl}}(P, 15), \quad \text{(S25)}$$

$$k_c^{\mathrm{monoalkyl}}(P, 60) = q(P) k_c^{\mathrm{diacyl}}(P, 60). \quad \text{(S26)}$$

The predicted osmotic pressure for each row was then

$$\pi = g k_c(P, \mathrm{backbone}, [\mathrm{PEG}]). \quad \text{(S27)}$$

Parameters were estimated by nonlinear weighted least squares.

This formulation yields PEG-specific 0-bar moduli

$$k_c^{\text{diacyl,15}} = a_{15}, k_c^{\text{diacyl,60}} = a_{60}, \quad \text{(S28)}$$
$$k_c^{\text{monoalkyl,15}} = q_0 a_{15}, k_c^{\text{monoalkyl,60}} = q_0 a_{60}. \quad \text{(S29)}$$

To report a single pair of 0-bar moduli for diacyl and monoalkyl POPE, we pooled the two diacyl estimates across 15 and 60% w/v PEG by generalized least squares using the fitted covariance matrix, and obtained the corresponding monoalkyl value by multiplying that pooled diacyl estimate by $q_0$. Fractional softening at 0 bar was reported as

$$100 \times \frac{k_{c,\text{DAG}} - k_{c,\text{AEG}}}{k_{c,\text{DAG}}} = 100(1 - q_0). \quad \text{(S30)}$$

Because the absolute moduli depend somewhat on model form, the monoalkyl softening factor is the most robust inference from this analysis; the pooled 0-bar $k_c$ values should be treated as model-estimated quantities.

Measurement of PC bilayer bending moduli using micropipette aspiration and optical tweezers

***GUV electroformation***

Powdered phospholipids were purchased from Avanti Polar Lipids and dissolved in HPLC-grade chloroform (Sigma Aldrich). For the vesicle formation, POPC (850457) or C16-18:1 PC (878112) was mixed with 4.995 mol% DSPE-PEG2000-Amine (880128), 0.005 mol% DSPE-PEG2000-Biotin (880129), and 0.5 mol% Liss Rhodamine DOPE (810150) to a total lipid concentration of 2.5mM. PEGylated lipids were added to prevent non-specific interactions (27). Using a Hamilton syringe with a bevelled needle, 6 µL solution of the lipid mixture was gently spread over a plasma-treated (Diener, 40% power $O_2$ plasma for 3 minutes) ITO-covered glass slide (Nanion Technologies) and dried under an argon stream. Subsequently, the slide was desiccated under mild vacuum for at least 2 hr before the electroformation.

The dried lipid cake was rehydrated with 170 mOsm $L^{-1}$ sucrose solution (~170 mM), and GUVs were formed using the Vesicle Prep Pro electroformation machine (Nanion Technologies). The electroformation voltage and frequency were increased stepwise to 5 V and 50 Hz, and maintained for 90 minutes at 55 °C, followed by a gradual reduction of the frequency to 5 Hz across an additional 50 minutes (28). The vesicles were stored at 4 °C within 24 hours.

***Optical Tweezers setup for bending rigidity measurements***

The experiments were performed using a C-trap confocal fluorescence optical tweezers setup (Lumicks) based on an inverted microscope equipped with a water-immersion objective (NA 1.2) and a condenser top lens (NA 1.4). The optical trap is generated by a 10 W 1064 nm laser. The displacement of the optically trapped beads from the center was measured using back-focal-plane interferometry of the condenser lens with a position-sensitive detector and converted into a force signal. The samples were illuminated with a bright-field 850 nm LED and imaged in transmission using a complementary metal-oxide-semiconductor (CMOS) camera.

***Confocal fluorescence microscopy***

The C-trap includes three fiber-coupled excitation lasers at 488 nm, 561 nm, and 638 nm. Scanning is performed with a fast-tip/tilt piezoelectric mirror. For confocal detection, the emitted fluorescence was descanned, separated from the excitation by a dichroic mirror, and filtered using emission filters (blue: 500–550 nm; green: 575–625 nm; red: 650–750 nm). Photons were counted by using a fiber-coupled single-photon counting module. The multimode fibers serve as pinholes, providing background rejection. For Rhodamine excitation, the 561 nm laser was used at 1% power (10.86 µW).

***Sample manipulation***

The experimental chamber consisted of a PDMS wall (prepared using the 184-silicone elastomer kit, SYLGARD™) mounted on a clean precision cover glass (CG15KH1, THORLABS) and placed on an automated XY-stage. GUVs and streptavidin polystyrene microparticles ('beads', Spherotech Inc.) of 3.15 μm diameter were added to the chamber and diluted with a degassed and filtered 165 mM glucose solution containing 1 mM NaCl to ensure biotin-streptavidin binding.

A micropipette aspiration setup, including a micromanipulator (uMp-3, Sensapex) holding a capillary of 5 μm diameter (VICbI-5-10-1-55, BioMedical Instruments) connected to a microfluidic flow controller (EZ-25; Fluigent), was integrated into our optical tweezers instrument. To reduce GUV binding to the capillary, the capillaries were passivated in DDW containing 2% w/v BSA (A7906, Sigma). The experimental chamber was intentionally kept unpassivated due to an observed softening of the GUVs when working with BSA or β-casein in the solution. By controlling the aspiration pressure, the membrane tension on the GUV was modified according to (29):

$$\gamma_{\mathrm{asp}} = \frac{\Delta P \cdot R_{\mathrm{pip}}}{2(1 - R_{\mathrm{pip}}/R_{\mathrm{Ve}})}, \tag{S31}$$

where $\gamma_{\mathrm{asp}}$ is the aspiration tension, $\Delta P$ is the micropipette suction pressure, $R_{\mathrm{Ve}}$ is the vesicle radius, and $R_{\mathrm{pip}}$ is the micropipette radius. Before each experiment, the zero-suction pressure was found by aspirating a bead into the pipette and reducing the suction pressure until the bead stopped moving.

***Diacyl and monoalkyl 16:0-18:1 PC bending rigidity measurements***
Following the pressure calibration, a vesicle was aspirated into the micropipette and held at a ~1.5 mbar aspiration pressure for a minute to validate proper aspiration and check for unilamellarity of the membrane. Then, a streptavidin-coated bead was trapped by an optical trap and held at a 5% laser power (~0.5 W). The trap stiffness was calibrated using a passive power-spectrum method. The laser power was kept low to prevent excessive fluid flow near the optical trap and to prevent the potential trapping of membrane debris in the sample.

The trapped bead was brought into contact with the aspirated GUV; following the binding of the bead, a membrane tube ('tether') was pulled, and the tether pulling force was monitored (Fig. S3). Next, the GUV tension was adjusted stepwise by changing the aspiration pressure. After each step, we allowed the pulling force to equilibrate before measuring the force. The direction of the tension change was altered between GUVs, and we performed bi-directional measurements to validate that this had no effect on the results. The bending rigidity was evaluated based on the relation derived from the Helfrich model (30):

$$f = 2\pi\sqrt{2K_c \cdot \gamma_{\mathrm{asp}}} + C, \tag{S32}$$

where $K_c$ is the bilayer bending modulus and $C$ is a parameter dependent on the bilayer's spontaneous curvature. We consider $C$ to be constant (31), yet the value might differ across GUVs, as electroformation results in slight variations between vesicles.

<u>Measurement of PE chain melting temperatures by DSC</u>

Each multilamellar vesicle (MLV) sample was prepared by drying a $CHCl_3$ solution of 3 mg lipid on the wall of a 4 mL amber borosilicate vial (Wheaton) under a stream of $N_2$, then further drying the resultant film under high vacuum for 30 min at room temperature. Films were hydrated in 1.50 mL 20 mM HEPES + 2 mM EDTA, pH 7.4, which was degassed by sonication under high vacuum and preheated to 37°C. After the buffer was added, vials were purged with $N_2$ and tumbled at 20 RPM, 30 min at 37 °C, then freeze-thawed 3 times using dry ice and a 37°C bath.

One milliliter of each MLV preparation was transferred to a 96 x 2 mL well plate, which was flushed with $N_2$, covered, and loaded into the autosampler of a nano DSC (TA Instruments). The autosampler compartment was held at 10 °C. $L_\beta \rightarrow L_\alpha$ transition temperatures ($T_m$) were resolved in heating scans from 0-100 °C at 1 °C/min. The less enthalpic $L_\alpha \rightarrow H_{II}$ phase transition (32) was not apparent, however a strong exotherm was observed in plasmenyl PE at 31.5 °C that may correspond to decomposition of the alkenyl ether linkage.

Measurement of C-laurdan GP across pressure-temperature space

Large unilamellar vesicles (LUVs) used for high-pressure fluorescence spectroscopy (HPFS) were derived from 0.50 mL aliquots of the MLVs prepared for DSC (above). C-laurdan probe (TOCRIS) was added to each aliquot (0.5 µL of 5 mM stock solution in DMSO for ~1:800 probe:lipid) and the MLVs were passed 15 times through an 0.1 µm polycarbonate track-etch filter (Whatman) using a handheld mini extruder (Avanti) at 37°C. Total volume of the LUV suspension was brought to approximately 1.25 mL with HEPES-EDTA buffer and the LUVs were then transferred to the round quartz cuvette of the high-pressure fluorescence cell (ISS).

The cuvette was installed in the high-pressure cell mounted in a fluorospectrophotometer (Cary Eclipse). Fluorescence intensities at ex/em 340/440 and 340/490 nm were recorded every 30 s. The high-pressure cell traversed a T-P grid using a pattern described previously (10), but with 5 °C T-steps from 0-60 °C (PCs) or 20-80 °C (PEs) and 100 bar P-steps from 0-1000 bar gauge pressure. Temperature was controlled using a recirculating bath (NESLAB RTE-111) with external Pt100 thermistor (Omega Engineering) and pressure was controlled using a high-pressure syringe pump (Teledyne SyriXus 65x) filled with distilled water. The experiment was coordinated using a custom Python script (https://github.com/octopode/spectackler) and continuously recorded T, P, and fluorescence data were co-registered by timestamp.

Atomistic molecular dynamics simulations and analysis

***System construction, equilibration, and production***

Classical molecular dynamics (MD) simulations were performed for 5 PE lipids, representing the different glycerol backbone chemistries considered in this study. With the aim of understanding the unique effects of each backbone, we simulated single-component membranes composed of 18:0/18:1 plasmenyl, 18:0/18:1 diacyl, 16:0/18:1 dialkyl, 16:0/18:1 monoalkyl, and 16:0/18:1 diacyl glycerol backbone lipids. We worked with two membrane sizes for each lipid type. The smaller membranes were roughly 10×10 nm$^2$, with between 170 and 200 lipids per leaflet; the larger membranes were 20×20 nm$^2$, with between 670 and 700 lipids per leaflet. Initial configurations for each system were generated with the CHARMMGUI web server (33–36). All simulations were performed using NAMD version 3.0 (37), lipids and ions were modeled with the CHARMM36 force field (38) and water was modeled using the TIP3P model (39).

Each system was equilibrated and brought to the desired temperature and pressure using a series of seven minimization and heating steps numbered i-vii in the following. During equilibration, restraints were applied to maintain the double bonds and glycerol geometries, with an initial restraint force constant of 500 kcal/mol. Restraints were also implemented to maintain a planar, bilayer structure during equilibration, using the colvars module to restrain the positions of the phosphates along the membrane normal with a 5 kcal/(mol·Å²) force constant. (i) The initial configuration was minimized for 10,000 steps using steepest descent. Steps ii through iii were performed under conditions of constant particle number, volume, and temperature. (ii) 125,000

Langevin dynamics steps with a timestep of 1 fs; (iii) 125,000 steps with a timestep of 1 fs with the strength of the chemical geometry restraints reduced to 200 kcal/mol. Planar restraints were kept constant. (iv) 125,000 Langevin dynamics steps with a timestep of 1 fs with the strength of the chemical geometry restraints reduced to 100 kcal/mol and the force constant for the planar restraint reduced to 2 kcal/(mol·$Å^2$), under conditions of constant particle number, pressure, and temperature (NPT) with the pressure controlled using the Langevin piston method (40, 41). The piston was configured with a period of 50 fs and a piston decay of 25 fs. All subsequent stages used the same NPT protocol. (v) 250,000 steps of Langevin dynamics were run with a timestep of 2 fs. The strength of the chemical geometry restraints was maintained at 100 kcal/mol, while the force constant for the planar restraint was reduced to 1 kcal/(mol·$Å^2$). (vi) Integration parameters from the previous equilibration stage were maintained (250,000 steps with a timestep of 2 fs). The strength of the chemical geometry restraints was reduced to 50 kcal/mol, and the force constant for the planar restraint was reduced to 0.2 kcal/(mol·$Å^2$). (vii) 250,000 steps of NPT dynamics were run with a timestep of 2 fs with all restraints removed. Throughout the seven equilibration stages, velocities were reassigned every 500 steps, and hydrogen positions were constrained by SHAKE (42).

After the equilibration process was completed, production simulations were run for at least 500 ns, in the isothermal-isobaric (NPT) ensemble. Temperature was held constant at 35 °C using a Langevin thermostat (43) with a damping coefficient of 1 $ps^{-1}$, and pressure was coupled independently in the directions normal and parallel to the interface. The pressure was controlled using the Langevin Piston method (with 50 fs and 25 fs for the period and piston decay, respectively), which allows fluctuations in the cell volume (40). A switching function was applied for long-range dispersion, with 10-12 Å for the truncated cutoff; the SHAKE algorithm was implemented with a tolerance of $10^{-8}$, and Particle Mesh Ewald (PME) was used to compute long-range electrostatics at every timestep (44) using a maximum grid spacing of 1 Å and interpolation order 6. Production simulations were analyzed after 75 ns to ensure the system was fully equilibrated by checking the relaxation of the system area. Then, each simulation was post-processed by unwrapping, centering on the membrane's time-averaged geometric center, and rewrapping the membrane within its periodic boundary conditions. The qwrap tool in VMD was used for the wrapping and unwrapping processes (45).

***Molecular fiducials and the Luzzati plane***

For each of the lipid types, using the smaller MD-simulated membrane system, we extracted the average transbilayer location of the phosphate ($PO_4$) group, the sn-1 and sn-2 radyl carbon alpha to the glycerol oxygen ($C_{O\text{-}\alpha}$), and the Luzzati plane. The lipid atom locations were computed by fitting the electron density profiles of the corresponding chemical groups to gaussian distributions; the reported average is the mean of the gaussian fit. The electron density profiles were obtained from histograms of the transbilayer positions of the selected atoms, using atomic coordinates extracted with MDtraj (46). The histogram was weighted by the corresponding frame cell volume to account for fluctuations in the simulation volume. The optimal number of histogram bins was estimated using Knuth's rule (47) and the histogram was normalized. The Luzzati plane corresponds to the position where the water density equals half of the bulk water value. The water electron density was computed using the water oxygen atom, as it provides a representative position of the water molecule's geometric center. The half bulk water value was determined by dividing the maximum water density by 2. To find the corresponding transbilayer location, linear interpolation was performed around the half bulk water value to avoid errors arising from bin discretization. The error in the Luzzati plane location was calculated by bootstrapping with 500 repetitions to ensure convergence.

As a comparison to the Luzzati plane, we also calculated the Gibbs dividing surface: the location where the integral of the water electron density reaches one-half of its maximum value, with the upper limit being the location at which the water density reaches its bulk value (48, 49). The water electron density was computed using the water oxygen atom, as it provides a representative position of the water molecule's geometric center. The upper limit for the integral was determined as the location at which the water density reaches its bulk value, the latter determined by averaging the water density over a region distant from the membrane. The precise location of the Gibbs surface value was located by linear interpolation to avoid errors arising from bin discretization. The error in the Gibbs surface location was calculated by dividing the trajectory into non-overlapping blocks, locating the Gibbs surface for each block, then computing the standard error over the blocks (50). This was then repeated for longer and longer blocks until convergence of the error estimate was obtained. Compared to the Luzzati plane, the Gibbs surface was ~0.3 nm closer to the bilayer center across all systems (Figure S5C).

***Bending modulus and neutral plane location***

The transverse curvature bias (TCB) method (51) provides a way to estimate the bending modulus by exploiting the molecular structure of the bilayer. When a bilayer bends, the two leaflets experience opposite lateral area strains: one region is locally expanded while the other is compressed. Within each leaflet, there exists a neutral plane at which bending does not produce local area expansion or compression. Lipid atoms located away from this neutral plane therefore sample a curvature that is systematically biased by their transverse position within the leaflet. In the TCB framework, auxiliary surfaces are defined at different distances $z$ from the neutral plane, and the average curvature sampled at those positions is measured. The key result is that the expected curvature bias is independent of the wavevector magnitude $\left\langle \bar{c}_{\vec{q}} \right\rangle$ and varies linearly with $z$:

$$\left\langle \bar{c}_{\vec{q}} \right\rangle = \frac{-2zk_BT}{L^2k_c} \quad \text{(S33)}$$

Thus, the slope of $\left\langle \bar{c}_{\vec{q}} \right\rangle$ as a function of the atomic position relative to the neutral plane is inversely proportional to $k_c$, providing an estimate of the bilayer bending modulus. Here, $k_BT$ is the thermal energy and $L^2$ is the mean area of the membrane.

Using the larger MD-simulated membrane system for each lipid type, TCB calculations were performed using the *MembraneAnalysis.jl* library in Julia (52). The library maps lipid positions onto a discretized membrane surface and computes the mean curvature spectrum $\left\langle \bar{c}_{\vec{q}} \right\rangle$. For this calculation, the average transverse position $z$ of the lipid heavy atoms is determined relative to the membrane neutral-surface reference, and the mean sampled curvature associated with those atomic positions is calculated. The sampled curvature field is then Fourier-transformed to obtain $\left\langle \bar{c}_{\vec{q}} \right\rangle$ as a function of the heavy-atom distance $z$.

***First moment and $c_0$ calculation***

For small bending deformations that are well-described by the Helfrich Hamiltonian, the product of $c_0$ and $k_c$ can be computed numerically from all-atom simulations of bilayers, by finding the first moment of the lateral pressure profile ($\pi(z)$):

$$-k_c c_0 = \int_{-\infty}^{\infty} z\pi(z)dz, \tag{S34}$$

where $z$ is the distance measured along the bilayer normal, with $z = 0$ at the center of the membrane.

The pressure profile was obtained for each simulated system by post-processing the production simulations. Each simulation snapshot (recorded every 100 ps) was used as an initial configuration for pressure profile calculation by assigning randomized velocities, running a short equilibrium simulation (1 ps), then computing the pressure profile using a modified version of NAMD that fixed a bug in the pressure profile routine (available at https://github.com/alexsodt/namd_profile_patch.git). The calculations employed the Harasima contour, with electrostatic interactions computed using PME as in the production run. For each calculation, the membrane was divided into 250 slabs along the bilayer normal. The lateral pressure in each slab was then used to determine $\pi(z)$, which was fitted with cubic splines to first interpolate the data and then to numerically compute the first moment (10).

***Lateral diffusion and area per lipid***

Lateral diffusion coefficients ($D_T$) were computed from the in-plane mean square displacement (MSD) of the phosphate group (53). Before computing the MSD the trajectories were "unwrapped" to remove discontinuities arising from lipids crossing periodic boundaries, followed by removal of the overall center of mass motion. The squared displacement was then computed for all lipids, and averaged over time and over individual lipids as implemented in the MDAnalysis Python library (54, 55). The diffusion coefficient was then obtained from the slope of a linear fit to the membrane MSD as a function of lag time, where the slope is equal to $4{*}D_T$. The bootstrapping method was used to estimate the errors in the diffusion coefficients, with 600 repetitions to ensure convergence. Because diffusion in a periodic membrane simulation depends strongly on box size (a finite-size hydrodynamic effect (56)), we evaluated the diffusion constant for both the 10 × 10 nm and 20 × 20 nm bilayers and compared the diffusion constants across the different lipid chemistries within each system size; the ordering of the diffusion constants was preserved across both sizes. Area per lipid (APL) was computed as the mean lateral area of the simulation cell divided by the number of lipids per leaflet.

Decomposition of energetic contributions to the $H_{II}$–$L_\alpha$ phase transition

Mechanical free-energy densities around the $H_{II}$→$L_\alpha$ transition were decomposed following the framework of Kirk et al., in which equality of phase free energies at coexistence gives

$$\Delta f_{\mathrm{IPF}}^{H_{\mathrm{II}}} + \Delta f_{\mathrm{CES}}^{H_{\mathrm{II}}} = \Delta f_{\mathrm{CES}}^{L_\alpha} \tag{S35}$$

Curvature elastic strain (CES) was calculated from the Helfrich form

$$\Delta f_{\mathrm{CES}} = k_c(c - c_0)^2 \tag{S36}$$

with $c_0$ the intrinsic monolayer curvature and $c$ the realized monolayer curvature. In the lamellar phase, mean monolayer curvature was taken as zero, such that

$$\Delta f_{\mathrm{CES}}^{L_\alpha} = k_c c_0^2 \tag{S37}$$

whereas in $H_{II}$, nonzero $c$ must be considered per Eq. S35. The interstitial packing frustration (ipf) contribution in $H_{II}$ was then obtained by difference at the $H_{II}$–$L_\alpha$ phase boundary:

$$\Delta f_{\mathrm{IPF}}^{H_{\mathrm{II}}} = \Delta f_{\mathrm{CES}}^{L_\alpha} - \Delta f_{\mathrm{CES}}^{H_{\mathrm{II}}}. \quad \text{(S38)}$$

Transition pressures were taken from the fitted barotropic phase boundaries, and uncertainty was propagated by bootstrap sampling of transition pressures ($P_{\mathrm{trans}} \sim \mathcal{N}(\hat{P}, \mathrm{SEM})$), curvature-model predictions for both relaxed ($c_0$) and unrelaxed ($c$) states, and measured bending moduli $k_c$ ($k_c \sim \mathcal{N}(\hat{k}_c, \mathrm{SEM})$). For $\Delta f_{\mathrm{IPF}}^{H_{\mathrm{II}}}$, terms were evaluated at each lipid's sampled $H_{II} \rightarrow L_\alpha$ transition pressure; for $\Delta f_{\mathrm{CES}}^{L_\alpha}$, ester-ether differences were evaluated at the midpoint between the two corresponding transition pressures so that both lipids were compared at a common reference pressure. Bootstrap distributions were summarized by their means and confidence intervals to obtain the reported $\Delta\Delta f$ values for structural substitutions.

### Extended Discussion

Role of interlamellar interactions

The free energy associated with dehydrating membrane surfaces (i.e. work of dehydration) is known to be a major barrier to membrane fusion (57). Our Kirk-like thermodynamic model of the $H_{II} \rightarrow L_\alpha$ transition does not explicitly include hydration terms; however, $f_{\mathrm{CES}}$ implicitly captures dehydration work associated with changes in headgroup packing during monolayer bending. In the $L_\alpha$ phase, free energy is additionally influenced by short-range hydration repulsion between membrane surfaces and attractive van der Waals interactions between adjacent bilayers, both of which depend on the interlamellar water gap $d_w$. Under the excess-water conditions used in our phase-transition experiments, the net interlamellar pressure is expected to vanish, bringing $d_w$ to an equilibrium value on the order of 5-10 Å (58, 59). Indeed, bilayer thicknesses of ~44 Å estimated from global fits to $L_\alpha$ SAXS profiles (60), when subtracted from measured repeat spacings of 49-62 Å, imply water gaps within this range. The free-energy well associated with bilayer adhesion at this equilibrium spacing (Fig. S7), which is about 3.0-3.6x$10^{-4}$ $k_BT/\text{Å}^2$ in POPE, provides an upper bound on the magnitude of interlamellar interaction free energy density (29). This value is an order of magnitude smaller than the mechanical free energy differences associated with backbone structure that we resolve here; we therefore omit explicit interlamellar interaction terms from $f_{L_\alpha}$.

Catalysis of membrane fusion by ether lipids

The similar lipid mixing rates across plasmenyl, dialkyl ether, and monoalkyl ether PE compositions, despite their distinct intrinsic curvatures, underscore that the kinetics of hemifusion cannot be predicted from $c_0$ alone. Elastic models of the classical fusion pathway estimate the free energy of fusion intermediates using multiple parameters including spontaneous curvature, the bending modulus, and the Gaussian modulus (61). These variables contribute to area stretch, lipid splay, and saddle splay that affect the energy barrier of hemifusion. Current experimental methods are unable to measure Gaussian curvature, but models estimate that it contributes on the order of 100 $k_BT$ to stalk and fusion pore energies (62, 63). Additionally, changes in lipid tilt and local variation in the Gaussian modulus can alter the fusion energy landscape (57, 64). Liposomes enriched in PE have been proposed to directly transition from a hemifusion stalk to the fusion pore, bypassing the formation of a hemifusion diaphragm (65). This suggests that alternate fusion pathways and energetic landscapes may exist that are governed by differences in elastic moduli. While the arrangement of lipids in fusion intermediates transitions through non-lamellar topologies similar to $H_{II}$ and $Q_{II}$, structures may differ based on lipid composition and environmental factors. For example, the propensity of the monoalkyl ether PE to form a $Q_{II}$ phase may produce a Gaussian curvature that largely offsets its lower $-c_0$ compared to the dialkyl ether PE or increased IPF compared to the plasmenyl PE. Compensating variations across elastic parameters along with effects that emerge locally may converge at similar fusion kinetics despite differences in intrinsic curvature.

### Supporting Figures

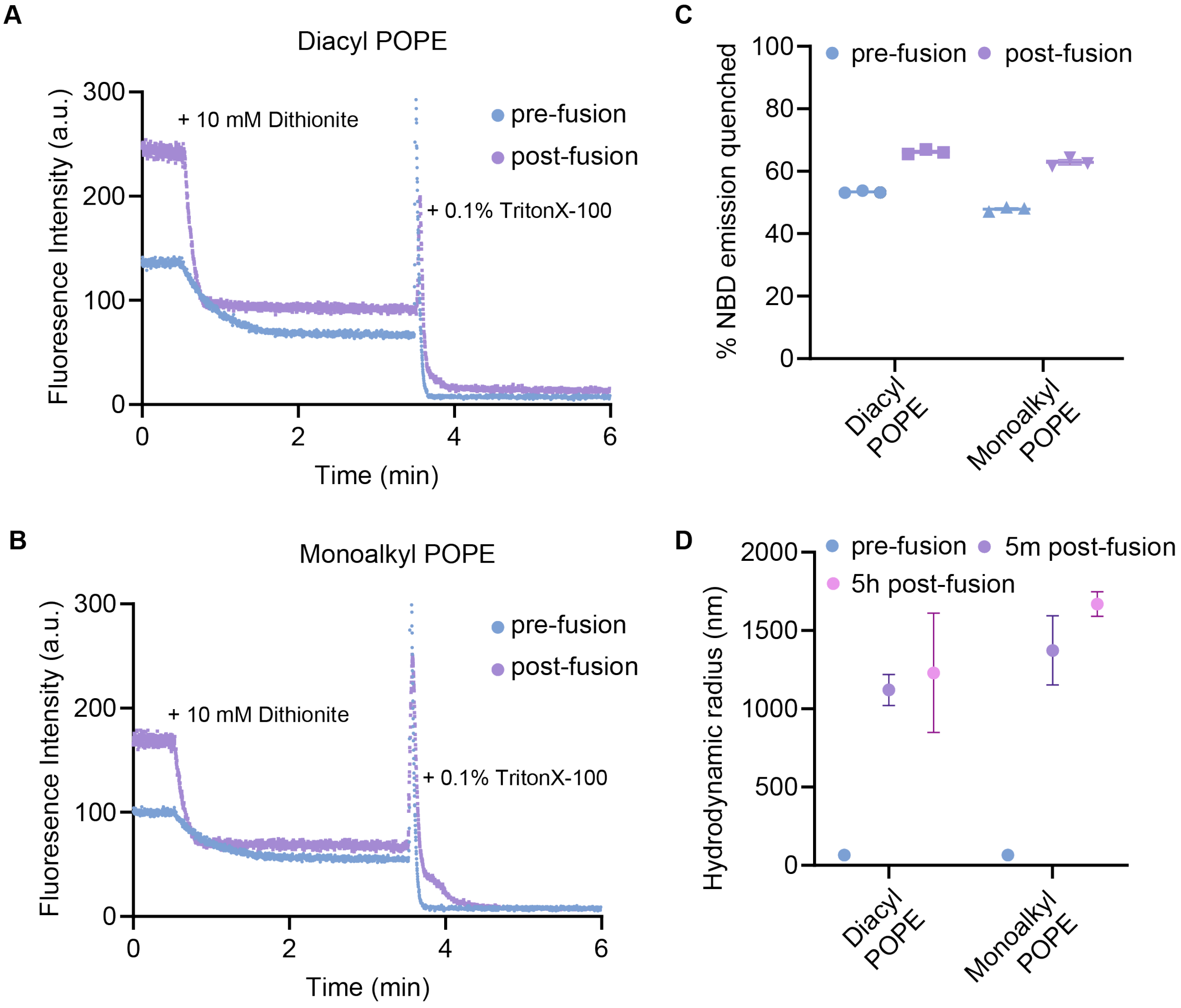


**Figure S1: Verification of vesicle unilamellarity and size for lipid mixing experiments.** (**A**) Representative NBD emission intensity in vesicles with 33:33:33:2 mol% diacyl POPE:POPC:POPS:NBD-PE following addition of 10 mM dithionite at 30 s and 0.1% Triton X-100 at 3.5 min. Dithionite quenches NBD-PE fluorescence but only on the outer leaflet in intact vesicles because of its low permeability. Triton X-100 addition dissolves the vesicles, allowing all NBD-PE to be quenched. (**B**) Representative NBD emission intensity in vesicles with 33:33:33:2 mol% monoalkyl POPE:POPC:POPS:NBD-PE following addition of 10 mM dithionite at 30 s and 0.1% Triton X-100 at 3.5 min. (**C**) Normalized percent NBD-PE emission quenched by dithionite from vesicles with diacyl and monoalkyl POPE before and after fusion by the addition of 10 mM $CaCl_2$. Data represents mean and SEM of triplicate measurements for each condition. Pre-fusion, ~50% of NBD emission is quenched for both linkage types, consistent with unilamellar vesicles. Post-fusion, there is a modest increase in dithionite quenching, consistent with a low level of leakage as previously observed for $Ca^{2+}$-mediated fusion of PS-containing vesicles (60) (**D**) Hydrodynamic radius (nm) measurements of vesicles by DLS before fusion was initiated by 10 mM $CaCl_2$, 5 minutes post-fusion, and 5 hours post-fusion. Before fusion, radii were 65 (diacyl) or 66 (monoalkyl) nm. Vesicle size increased post-fusion for both systems. Points represent mean and SEM of triplicate measurements for each condition.

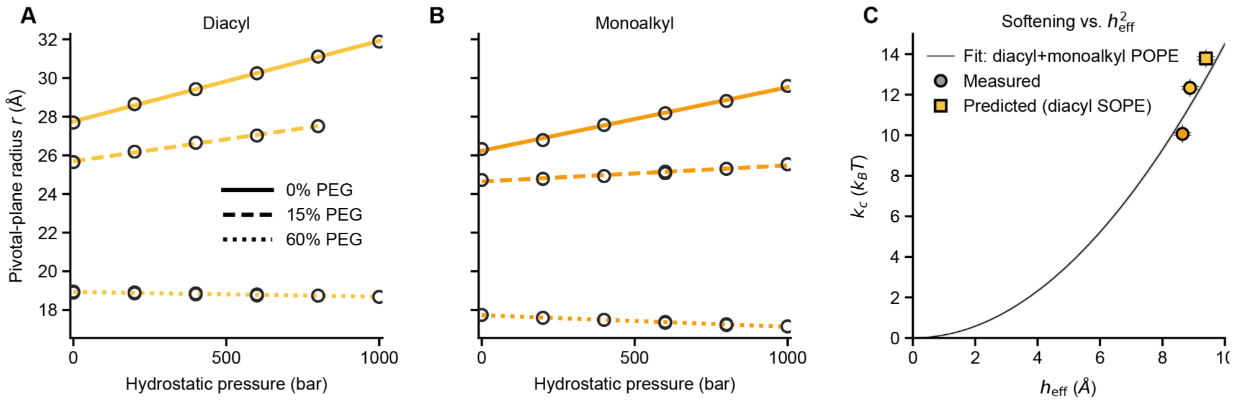


**Figure S2: Measurement of monolayer bending modulus for PEs.** (**A**) $H_{II}$ radius $r$ near the neutral plane as a function of osmotic and hydrostatic pressure for 16:0/18:1 diacyl PE. Osmotic pressure $\pi$ expels water from the $H_{II}$ tubes, making them smaller. (**B**) Measurements shown in **A**, but for monoalkyl PE. For a given level of $\pi$, monoalkyl monolayers bend more relative to their intrinsic radius, reflecting their lower bending rigidity. Divergence of the hydrostatic pressure trends reflects an apparent softening with hydrostatic pressure. (**C**) Softening by the monoalkyl backbone is stronger than predicted by elastic plate theory, where $k_c$ is expected to scale as the square of effective hydrocarbon thickness $h_{\text{eff}}$. The bending modulus for diacyl SOPE is predicted using this relation. Error intervals on the points and fitted parabola are ±SEM.

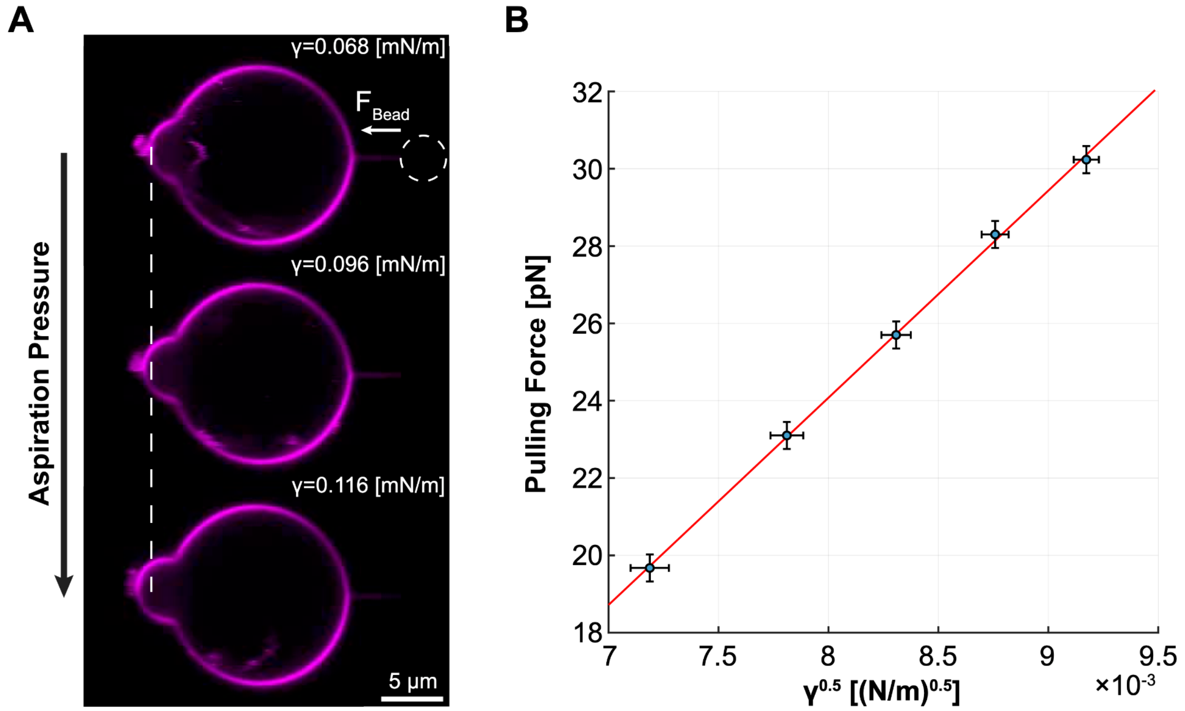


**Figure S3: Bending rigidity measurement in optical tweezers setup.** (**A**) Experimental setup – a tether is pulled from a micropipette aspirated GUV with an optically trapped streptavidin-coated bead (dashed circle), and the tether pulling force on the bead is measured. An increase in aspiration pressure pulls membrane area into the micropipette (the aspirated tongue, labelled with a dashed line, increases in length) and induces higher membrane tension, which subsequently increases tether pulling force. The fluorescence scans depict a single monoalkyl GUV, scale bar = 5μm. (**B**) Tether pulling force as a function of the square root of the membrane tension. The red line represents a linear fit, and the bending rigidity is evaluated from the slope, which equals $2\pi\sqrt(2K_c)$ (see Materials and Methods). The data represent a measurement from a single diacyl GUV.

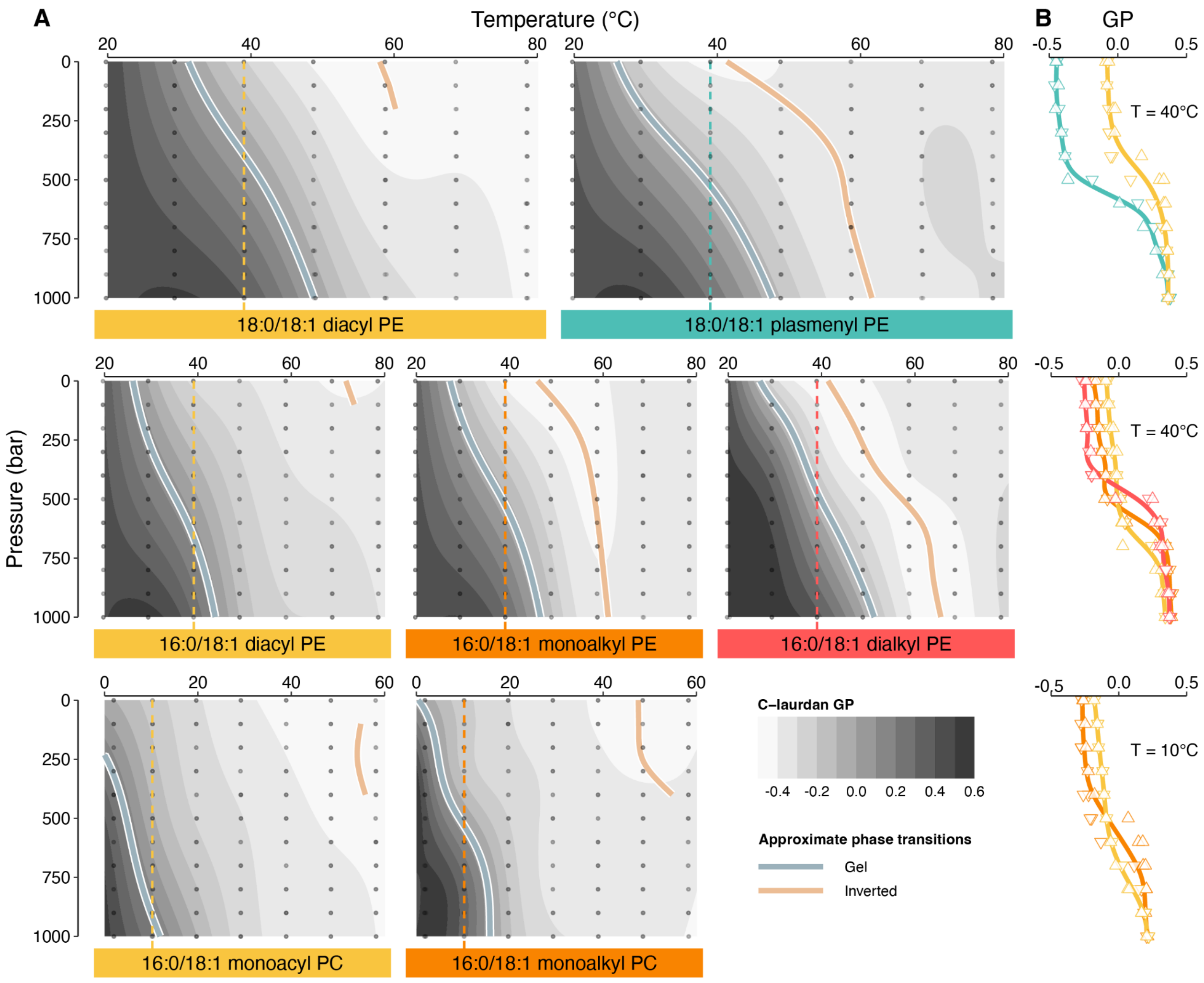


**Figure S4: C-laurdan fluorescence spectroscopy of backbone series** (**A**) Temperature-pressure landscapes of C-laurdan GP. Pressure is shown increasing downward and the grayscale contour interval is 0.1 GP units. Dashed vertical lines show the constant T-levels used to construct the P-slices shown in **B**. Light blue and salmon colored lines indicate approximate $L_\beta$-$L_\alpha$ and $L_\alpha$-$H_{II}$ boundaries, respectively. The former were determined from the inflection points of the P-slices; the latter follow ridges in the GP landscape. The proximity of these boundaries to each other illustrates the narrowing of the $L_\alpha$ stability region by ether-linked backbones. (**B**) Pressure slices of GP at constant temperatures chosen to bracket the $L_\alpha$-$L_\beta$ transition. Point shape indicates the P-sweep direction. The intersecting splines in the alkyl-PE and PC series show that more gel-prone ether-linked backbones exhibit lower GP in the $L_\alpha$ phase, due to increased water penetration that is not coupled to chain ordering. Plasmenyl PE shows a much larger GP difference from its diacyl analog, presumably due to the water-penetration effect of the ether in combination with the mild chain-disordering effect of the C=C bond.

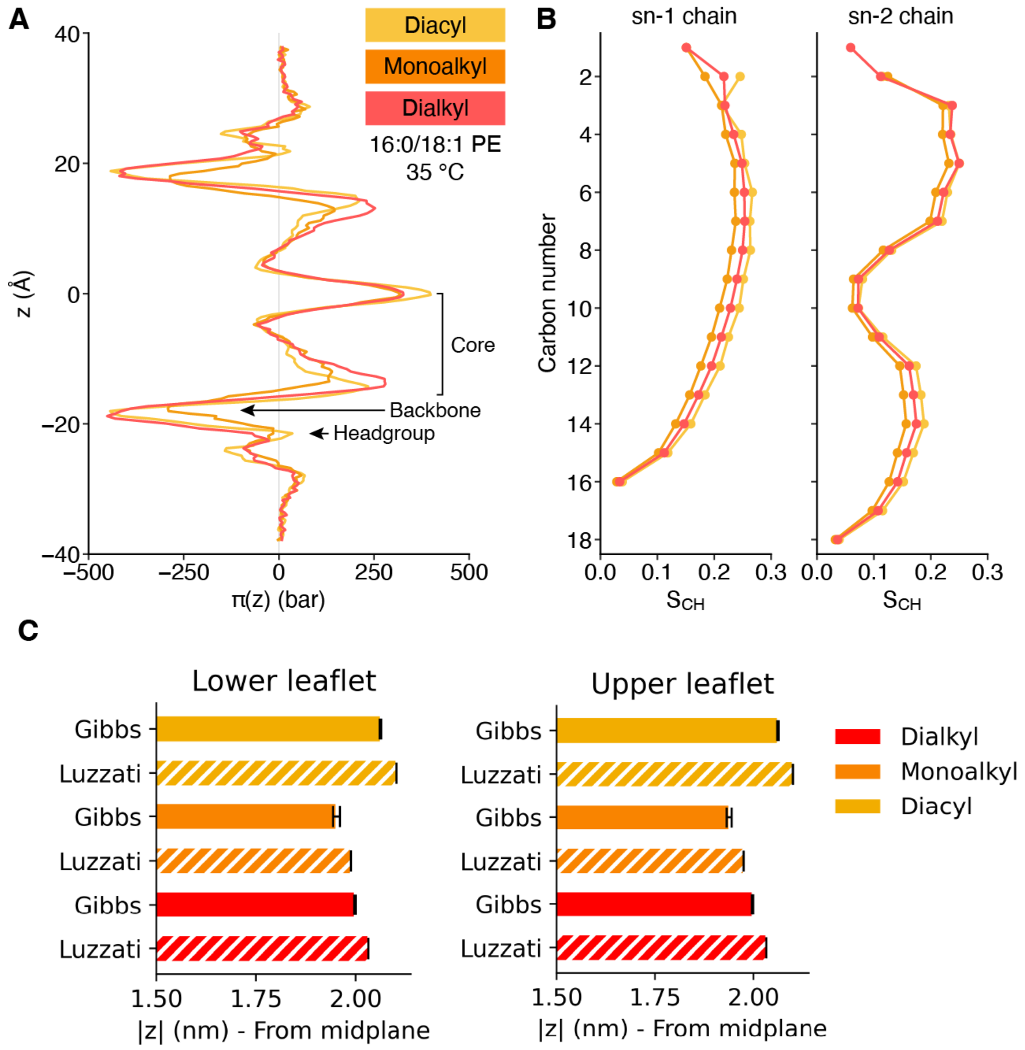


**Figure S5: Lateral pressure profiles, C-H order parameter profiles, and comparison of Luzzati plane and Gibbs surface from simulations of the alkyl linkage series** (**A**) Superimposed lateral pressure profiles at 35 °C and 1 bar hydrostatic pressure. The overall shape of these profiles reflects the PE lipids' nonbilayer tendency: the hydrophobic core exhibits high lateral pressure and the headgroup peaks are small. In the two more conical ether-linked species, the headgroup plane is under tension. (**B**) Superimposed profiles of the C-H order parameters for the carbons in each chain. With the exception of sn-1 carbon 2 (the O-β carbon), these order parameters are similar across the backbone series, reflecting the modest impact of alkyl linkages on chain order. (**C**) Differences between the calculated Luzzati plane and Gibbs surface – two measures of water penetration – for the series. Across both leaflets, the Gibbs surface reports a deeper penetration (closer to the midplane) of 0.03-0.04 nm, with the difference consistent across all series.

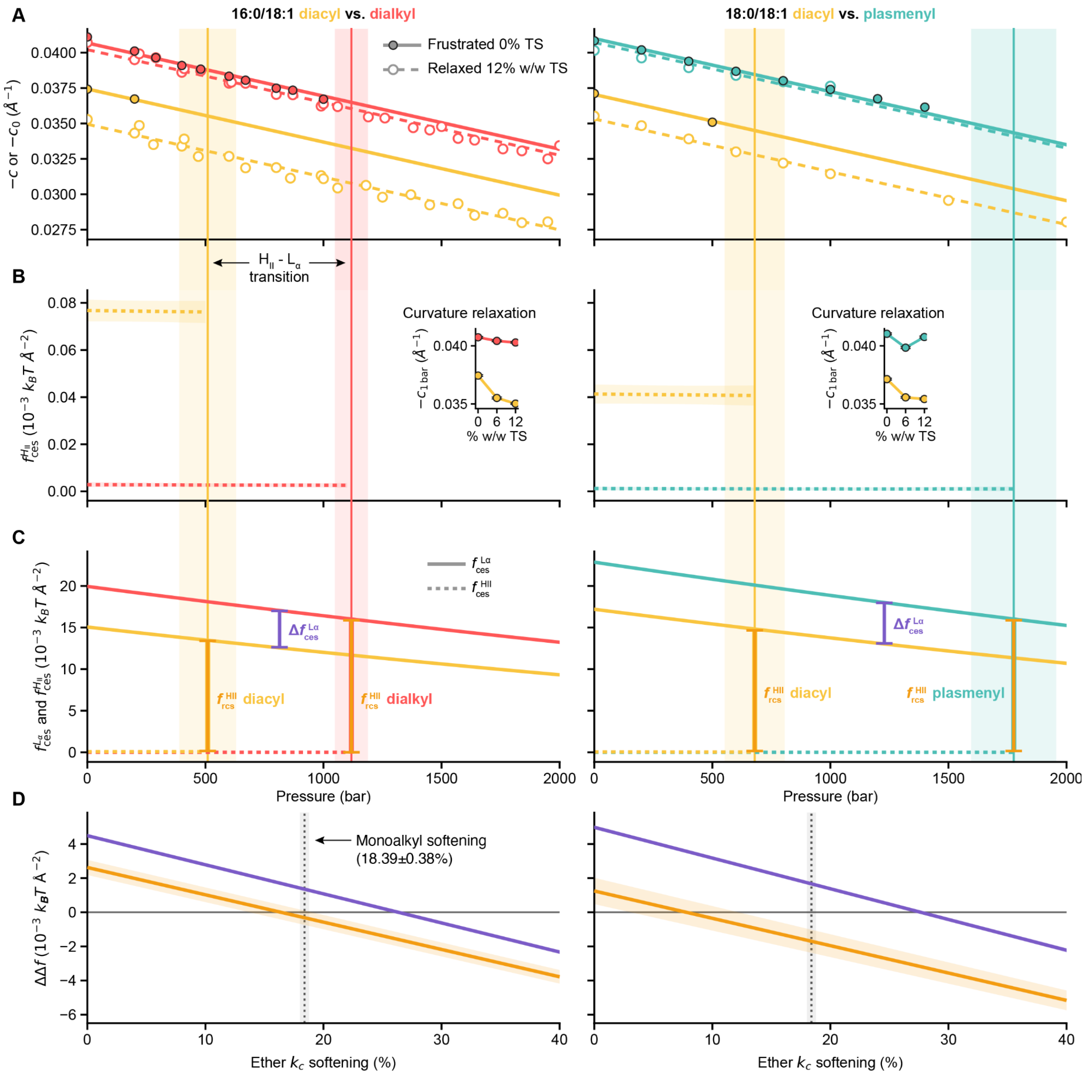


**Figure S6: Free-energy decomposition and derivation of $\Delta\Delta f_{\mathrm{mech}}$** (**A**) Frustrated and relaxed (= intrinsic) curvatures at the pivotal plane, overlaid with H$_{II}$–L$_α$ transition pressures for each lipid (vertical lines with ±SEM error bands) and predictions (solid and dashed lines) from the global linear model of curvature (see Methods), which was used to extrapolate frustrated curvature to the transition pressure. (**B**) CES energy density in the frustrated H$_{II}$ phase ($f_{\mathrm{CES}}^{H_{\mathrm{II}}}$), estimated using experimentally derived diacyl bending moduli (Fig. S2), the monoalkyl softening factor, and curvature relaxation by TS "filler" (insets). The global effect of pressure in our curvature model makes each constant. (**C**) CES energy density in L$_α$ ($f_{\mathrm{CES}}^{L\alpha}$) overlaid with the much smaller $f_{\mathrm{CES}}^{H_{\mathrm{II}}}$. $f_{\mathrm{CES}}^{L\alpha}$ was evaluated at the midpoint between phase transitions of the two lipids (purple vertical bar), while $f_{\mathrm{IPF}}^{H_{\mathrm{II}}}$ was calculated for each of the two lipids at their respective transitions (orange vertical bars) and $f_{\mathrm{IPF}}^{H_{\mathrm{II}}}$ was taken as the difference of these. (**D**) Free energy components $f_{\mathrm{CES}}^{L\alpha}$ and $f_{\mathrm{IPF}}^{H_{\mathrm{II}}}$ depend on the monolayer-softening effect of the backbone: a softer ether lipid reduces the destabilization of L$_α$ and strengthens the apparent contribution to IPF. The monoalkyl softening effect (dotted line) is used as the central scenario. The total effect on the mechanical free energy of phase transition, $\Delta\Delta f_{\mathrm{mech}}$, corresponds to the gap between the purple and orange lines and is nearly invariant with softening.

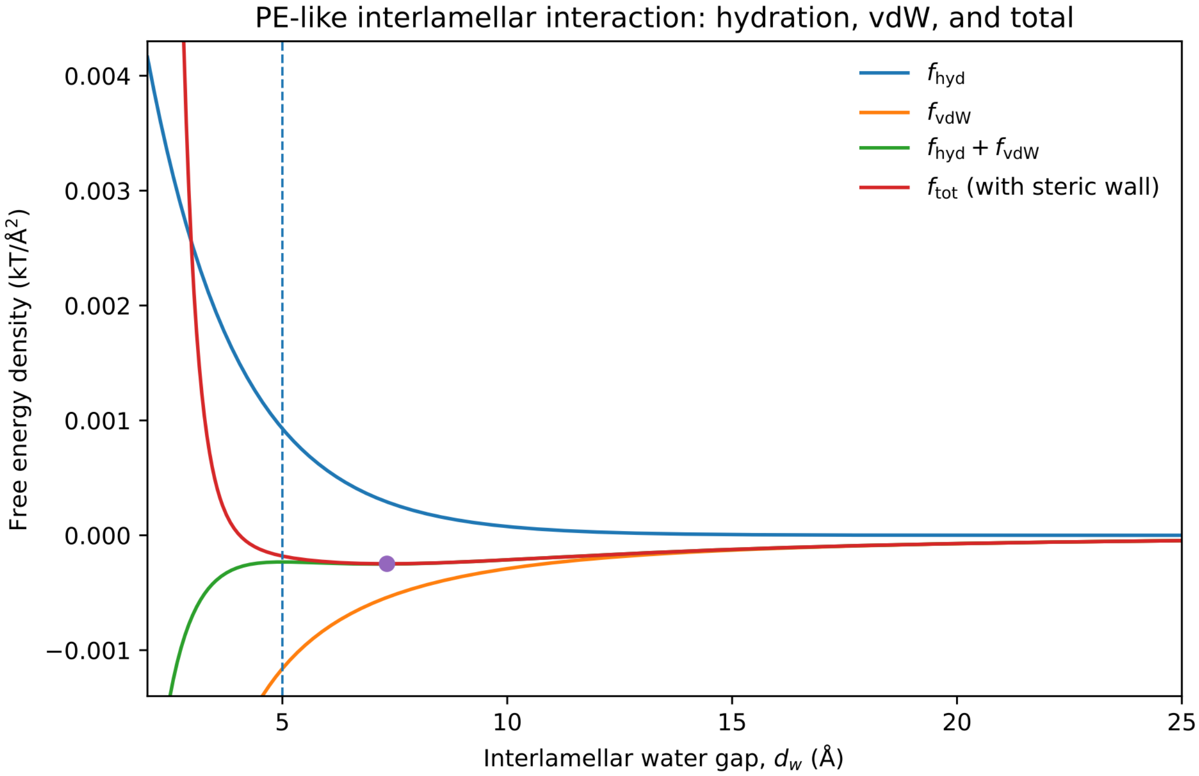


**Figure S7: Model of interlamellar interaction free energy for PE bilayers.** Hydration repulsion (blue) was modeled as $f_{\text{hyd}} = P_0 \lambda e^{-d_w/\lambda}$ and van der Waals attraction (orange) as $f_{\text{vdW}} = -A/(12\pi d_w^2)$, using representative values $\lambda = 2.0$ Å and $A = 4.5 \times 10^{-21}$ J. $P_0 = 5.66 \times 10^{-3}\ k_B T/\text{Å}^3$ was chosen such that the corresponding hydration and van der Waals pressures balance at $d_w = 5$ Å (dashed line), consistent with equilibrium water gaps reported for fully hydrated PE lamellae (58, 59). The sum of these terms is shown in green. To prevent unphysical collapse of the continuum van der Waals potential at very small separation, a short-range steric repulsion $f_{\text{steric}} = B/d_w^8$, with $B = 20\ k_B T \text{Å}^6$, was added to give $f_{\text{tot}}$ (red). The resulting interaction potential has a shallow minimum at $d_w \approx 7.3$ Å (purple point).

### Supporting Tables

**Table S1: Proportions of different backbone types under PE and PC headgroups across the animal tree of life.** Each backbone is quantified as mole fraction of its respective headgroup. Data are listed in the same order as they are plotted in Fig. 1C.

| Phylum | Species | Life stage | Tissue | Ref. | PC | | | PE | | |
|---|---|---|---|---|---|---|---|---|---|---|
| | | | | | Diacyl | Mono-alkyl | Plas-menyl | Diacyl | Mono-alkyl | Plas-menyl |
| Ctenophora | *Platyctenida* sp. T | Adult | Whole body | (10) | 0.874 | 0.059 | 0.067 | 0.269 | 0.053 | 0.678 |
| Ctenophora | *Bolinopsis vitrea* | Adult | Whole body | | 0.869 | 0.070 | 0.061 | 0.361 | 0.022 | 0.617 |
| Ctenophora | *Lampocteis cruentiventer* | Adult | Whole body | | 0.834 | 0.093 | 0.073 | 0.590 | 0.054 | 0.356 |
| Ctenophora | *Bolinopsis microptera* | Adult | Whole body | | 0.965 | 0.023 | 0.012 | 0.741 | 0.030 | 0.229 |
| Ctenophora | *Bolinopsis infundibulum* | Adult | Whole body | | 0.959 | 0.026 | 0.016 | 0.866 | 0.016 | 0.118 |
| Porifera | *Sphaerotylus borealis* | Adult | Whole body | (66) | 0.562 | 0.247 | 0.191 | 0.120 | 0.321 | 0.559 |
| Porifera | *Polyrnastia sp.* | Adult | Whole body | | 0.315 | 0.516 | 0.168 | 0.000 | 0.584 | 0.416 |
| Porifera | *Suberites domuncula* | Adult | Whole body | | 0.725 | 0.148 | 0.128 | 0.177 | 0.500 | 0.323 |
| Porifera | *Oscarella lobularis* | Adult | Whole body | | 0.685 | 0.176 | 0.139 | 0.231 | 0.458 | 0.311 |
| Porifera | *Suberites sp.* | Adult | Whole body | | 0.702 | 0.113 | 0.185 | 0.000 | 0.746 | 0.254 |
| Porifera | *Hymedesmia sp.* | Adult | Whole body | | 0.765 | 0.188 | 0.047 | 0.000 | 0.864 | 0.136 |
| Porifera | *Halichondria panicea* | Adult | Whole body | | 0.598 | 0.315 | 0.087 | 0.083 | 0.785 | 0.133 |
| Porifera | *Mycale lobata* | Adult | Whole body | | 0.767 | 0.233 | 0.000 | 0.352 | 0.526 | 0.122 |
| Porifera | *Haliclona cinerea* | Adult | Whole body | | 0.459 | 0.541 | 0.000 | 0.248 | 0.636 | 0.116 |
| Porifera | *Haliclona aqueductus* | Adult | Whole body | | 0.848 | 0.114 | 0.038 | 0.196 | 0.701 | 0.103 |
| Porifera | *Phakellia sp.* | Adult | Whole body | | 0.784 | 0.187 | 0.029 | 0.072 | 0.829 | 0.099 |
| Porifera | *Myxilla sp.* | Adult | Whole body | | 0.553 | 0.401 | 0.046 | 0.404 | 0.503 | 0.092 |
| Porifera | *Mycale sp.* | Adult | Whole body | | 0.558 | 0.276 | 0.166 | 0.206 | 0.705 | 0.089 |
| Porifera | *Halichondria sitiens* | Adult | Whole body | | 0.611 | 0.330 | 0.059 | 0.115 | 0.799 | 0.085 |
| Porifera | *Haliclona sp.* | Adult | Whole body | | 0.815 | 0.185 | 0.000 | 0.178 | 0.745 | 0.077 |
| Porifera | *Dysidea fragilis* | Adult | Whole body | | 0.851 | 0.142 | 0.007 | 0.344 | 0.580 | 0.076 |
| Porifera | *Plocamia ambigua* | Adult | Whole body | | 0.851 | 0.108 | 0.040 | 0.586 | 0.365 | 0.049 |
| Porifera | *Hymeniacidon gorbunovi* | Adult | Whole body | | 0.633 | 0.273 | 0.094 | 0.112 | 0.865 | 0.023 |
| Porifera | *Gellius augulatus* | Adult | Whole body | | 0.746 | 0.234 | 0.021 | 0.360 | 0.625 | 0.016 |
| Mollusca | *Donax trunculus* | Adult | Whole body | (67) | 0.604 | 0.347 | 0.049 | 0.109 | 0.066 | 0.825 |
| Mollusca | *Pteria aegyptia* | Adult | Whole body | | 0.674 | 0.278 | 0.048 | 0.118 | 0.078 | 0.804 |
| Mollusca | *Unio terminalis* | Adult | Whole body | | 0.663 | 0.286 | 0.051 | 0.143 | 0.069 | 0.788 |
| Mollusca | *Callista florida* | Adult | Whole body | | 0.638 | 0.305 | 0.057 | 0.114 | 0.113 | 0.773 |
| Mollusca | *Mactra corallina* | Adult | Whole body | | 0.550 | 0.388 | 0.062 | 0.147 | 0.088 | 0.765 |
| Mollusca | *Potanida littoralis* | Adult | Whole body | | 0.615 | 0.343 | 0.042 | 0.174 | 0.094 | 0.732 |
| Mollusca | *Corbicula fluminalis* | Adult | Whole body | | 0.661 | 0.312 | 0.027 | 0.183 | 0.103 | 0.714 |
| Mollusca | *Mytilus galloprovincialis* | Adult | Whole body | | 0.665 | 0.297 | 0.038 | 0.145 | 0.158 | 0.697 |
| Arthropoda | *Homarus gammarus* | Adult | Leg | (68) | 0.720 | 0.280 | 0.000 | 0.176 | 0.310 | 0.514 |
| Arthropoda | *Homarus gammarus* | Adult | Claw | | 0.729 | 0.230 | 0.041 | 0.289 | 0.330 | 0.381 |
| Arthropoda | *Chloridea virescens* | Larva | Whole body | (69) | 0.997 | 0.003 | 0.000 | 0.794 | 0.046 | 0.160 |
| Arthropoda | *Chloridea virescens* | Pupa | Whole body | | 1.000 | 0.000 | 0.000 | 0.933 | 0.015 | 0.052 |
| Arthropoda | *Chloridea virescens* | Embryo | Whole body | | 0.997 | 0.003 | 0.000 | 0.957 | 0.021 | 0.022 |
| Chordata | *Chacharhinus plumbeus* | Adult | White muscle | (70) | 0.797 | 0.059 | 0.144 | 0.410 | 0.042 | 0.548 |
| Chordata | *Prionace glauca* | Adult | White muscle | | 0.844 | 0.088 | 0.068 | 0.518 | 0.000 | 0.482 |
| Chordata | *Alopias vulpinus* | Adult | White muscle | | 0.945 | 0.026 | 0.029 | 0.605 | 0.031 | 0.364 |
| Chordata | *Lamna ditropis* | Adult | White muscle | | 0.864 | 0.072 | 0.064 | 0.563 | 0.099 | 0.338 |

| | | | | | | | | | | |
|---|---|---|---|---|---|---|---|---|---|---|
| Chordata | *Lamna ditropis* | Adult | Dark muscle | (71) | 0.731 | 0.147 | 0.121 | 0.700 | 0.053 | 0.247 |
| Chordata | *Mus musculus* | Adult | Lung | | 0.981 | 0.005 | 0.014 | 0.392 | 0.010 | 0.597 |
| Chordata | *Mus musculus* | Adult | Colon | | 0.959 | 0.021 | 0.020 | 0.434 | 0.023 | 0.543 |
| Chordata | *Mus musculus* | Adult | Cerebellum | | 0.989 | 0.003 | 0.008 | 0.463 | 0.004 | 0.532 |
| Chordata | *Mus musculus* | Adult | Cerebrum | | 0.988 | 0.003 | 0.010 | 0.496 | 0.005 | 0.499 |
| Chordata | *Mus musculus* | Adult | Spleen | | 0.941 | 0.054 | 0.005 | 0.575 | 0.038 | 0.388 |
| Chordata | *Mus musculus* | Adult | Ovary | | 0.983 | 0.011 | 0.006 | 0.596 | 0.095 | 0.309 |
| Chordata | *Mus musculus* | Adult | Heart | | 0.980 | 0.008 | 0.012 | 0.702 | 0.001 | 0.297 |
| Chordata | *Mus musculus* | Adult | Kidney | | 0.896 | 0.097 | 0.007 | 0.754 | 0.002 | 0.244 |
| Chordata | *Mus musculus* | Adult | Testis | | 0.964 | 0.029 | 0.007 | 0.489 | 0.269 | 0.242 |
| Chordata | *Mus musculus* | Adult | Skel. muscle | | 0.992 | 0.003 | 0.005 | 0.766 | 0.002 | 0.232 |
| Chordata | *Mus musculus* | Adult | Liver | | 0.997 | 0.002 | 0.001 | 0.981 | 0.002 | 0.017 |
| Chordata | *Homo sapiens* | Adult | Adipose | (72) | 0.836 | 0.056 | 0.108 | 0.262 | 0.007 | 0.731 |
| Chordata | *Homo sapiens* | Adult | Aortic valve | (73) | 0.938 | 0.028 | 0.034 | 0.267 | 0.004 | 0.730 |
| Chordata | *Homo sapiens* | Adult | Brain | (74) | 0.696 | 0.165 | 0.139 | 0.435 | 0.073 | 0.492 |

**Table S2: Univariate sensitivity analysis of literature-derived input parameters to the lipid volume model.** The effects of uncertainty in these parameters on curvatures and monolayer effective thicknesses are quantified here. Input parameters are defined in Methods. Extreme low- and high-end estimates for each input parameter are shown, followed by the resultant absolute and percent changes to the output value relative to the central estimates used throughout our analyses (see Methods). The input parameters are ranked by decreasing impact, based on the larger of the low/high percent effects. Shaded rows indicate input parameters whose uncertainty effects are below the order of magnitude of resolved differences in the output parameter.

**(A)** $c$ **of unhosted samples at 75 °C**. No parameters have effects ≥10%.

| Parameter | Units | Low | High | Low $\lvert\Delta c\rvert$ (Å$^{-1}$) | High $\lvert\Delta c\rvert$ (Å$^{-1}$) | Low $\lvert\%\Delta c\rvert$ | High $\lvert\%\Delta c\rvert$ | Impact rank |
|---|---|---|---|---|---|---|---|---|
| $R_{\mathrm{aniso}}$ | | 2 | 4 | 0.001 | 0.00129 | 3.66 | 4.89 | 1 |
| $\kappa_V$ | bar$^{-1}$ | 3.5×10$^{-5}$ | 5.5×10$^{-5}$ | 4.92×10$^{-4}$ | 5.8×10$^{-4}$ | 1.86 | 2.21 | 2 |
| $a_{p,0}^{\mathrm{PE}}$ | Å$^2$ | 62.2 | 66.2 | 4.45×10$^{-4}$ | 4.15×10$^{-4}$ | 1.08 | 1.01 | 3 |
| $\alpha_A$ | K$^{-1}$ | 0.002 | 0.003 | 3.63×10$^{-4}$ | 3.4×10$^{-4}$ | 1.05 | 0.984 | 4 |
| $a_{p,\mathrm{sat}}^{\mathrm{PE}}$ | Å$^2$ | 71.6 | 75.6 | 3.49×10$^{-4}$ | 3.28×10$^{-4}$ | 0.857 | 0.807 | 5 |
| $V_{\mathrm{HC}}^{\mathrm{DOPE}}$ | Å$^3$ | 835 | 875 | 3.33×10$^{-4}$ | 3.35×10$^{-4}$ | 0.809 | 0.813 | 6 |
| $\Delta V_{\mathrm{ether}}$ | Å$^3$ | -20 | -10 | 1.67×10$^{-4}$ | 1.67×10$^{-4}$ | 0.405 | 0.406 | 7 |
| $\alpha_V$ | K$^{-1}$ | 8×10$^{-4}$ | 0.0012 | 1.44×10$^{-4}$ | 1.44×10$^{-4}$ | 0.371 | 0.372 | 8 |
| $V_{\mathrm{TS}}$ | Å$^3$ | 630 | 710 | 3.9×10$^{-5}$ | 2.73×10$^{-5}$ | 0.121 | 0.0702 | 9 |
| $V_{\mathrm{CH_2}}$ | Å$^3$ | 26.5 | 28.7 | 3.55×10$^{-5}$ | 3.55×10$^{-5}$ | 0.0866 | 0.0866 | 10 |
| $V_{\mathrm{CH=CH}}$ | Å$^3$ | 39.2 | 43.2 | 3.34×10$^{-5}$ | 3.34×10$^{-5}$ | 0.0811 | 0.0811 | 11 |
| $\Delta a_p^{\mathrm{PC}}$ | Å$^2$ | 1 | 3 | 0 | 0 | 0 | 0 | 12 |

**(B)** $c_0$ **of guest lipids at 35 °C.** Shaded parameters have effects <10%.

| Parameter | Units | Low | High | Low $\lvert\Delta c_0\rvert$ (Å$^{-1}$) | High $\lvert\Delta c_0\rvert$ (Å$^{-1}$) | Low $\lvert\%\Delta c_0\rvert$ | High $\lvert\%\Delta c_0\rvert$ | Impact rank |
|---|---|---|---|---|---|---|---|---|
| $V_{\mathrm{TS}}$ | Å$^3$ | 630 | 710 | 2.49×10$^{-4}$ | 1.35×10$^{-4}$ | 33.4 | 5.77 | 1 |

| | | | | | | | | |
|---|---|---|---|---|---|---|---|---|
| $V_{\mathrm{HC}}^{\mathrm{DOPE}}$ | Å$^3$ | 835 | 875 | $2.96\times10^{-4}$ | $2.97\times10^{-4}$ | 21.9 | 23.2 | 2 |
| $R_{\mathrm{aniso}}$ | | 2 | 4 | $9.17\times10^{-5}$ | $2.54\times10^{-4}$ | 4.69 | 22 | 3 |
| $a_{p,\mathrm{sat}}^{\mathrm{PE}}$ | Å$^2$ | 71.6 | 75.6 | $2.36\times10^{-4}$ | $2.71\times10^{-4}$ | 16.1 | 7.79 | 4 |
| $\Delta a_p^{\mathrm{PC}}$ | Å$^2$ | 1 | 3 | $1.13\times10^{-4}$ | $1.13\times10^{-4}$ | 15.2 | 15.1 | 5 |
| $\kappa_V$ | bar$^{-1}$ | $3.5\times10^{-5}$ | $5.5\times10^{-5}$ | $7.67\times10^{-5}$ | $1.31\times10^{-4}$ | 6.43 | 12.6 | 6 |
| $a_{p,0}^{\mathrm{PE}}$ | Å$^2$ | 62.2 | 66.2 | $1.21\times10^{-4}$ | $1.25\times10^{-4}$ | 5.39 | 5.55 | 7 |
| $\alpha_A$ | K$^{-1}$ | 0.002 | 0.003 | $5.84\times10^{-5}$ | $5.72\times10^{-5}$ | 3.51 | 2.34 | 8 |
| $V_{\mathrm{CH_2}}$ | Å$^3$ | 26.5 | 28.7 | $2.23\times10^{-5}$ | $2.23\times10^{-5}$ | 2.67 | 2.67 | 9 |
| $V_{\mathrm{CH=CH}}$ | Å$^3$ | 39.2 | 43.2 | $2.15\times10^{-5}$ | $2.15\times10^{-5}$ | 2.43 | 2.43 | 10 |
| $\alpha_V$ | K$^{-1}$ | $8\times10^{-4}$ | 0.0012 | $2.48\times10^{-5}$ | $2.49\times10^{-5}$ | 1.13 | 1.25 | 11 |
| $\Delta V_{\mathrm{ether}}$ | Å$^3$ | -20 | -10 | $1.08\times10^{-4}$ | $1.08\times10^{-4}$ | 0.801 | 0.801 | 12 |

**(C)** $h_{\mathrm{eff}}$ **of unhosted samples at 75 °C.** Shaded parameters have effects <1%.

| Parameter | Units | Low | High | Low $\lvert\Delta h_{\mathrm{eff}}\rvert$ (Å) | High $\lvert\Delta h_{\mathrm{eff}}\rvert$ (Å) | Low $\lvert\%\Delta h_{\mathrm{eff}}\rvert$ | High $\lvert\%\Delta h_{\mathrm{eff}}\rvert$ | Impact rank |
|---|---|---|---|---|---|---|---|---|
| $R_{\mathrm{aniso}}$ | | 2 | 4 | 1.25 | 1.8 | 10.8 | 15.4 | 1 |
| $\kappa_V$ | bar$^{-1}$ | $3.5\times10^{-5}$ | $5.5\times10^{-5}$ | 0.625 | 0.821 | 5.37 | 7.05 | 2 |
| $\alpha_A$ | K$^{-1}$ | 0.002 | 0.003 | 0.35 | 0.31 | 3 | 2.66 | 3 |
| $a_{p,0}^{\mathrm{PE}}$ | Å$^2$ | 62.2 | 66.2 | 0.296 | 0.282 | 2.64 | 2.51 | 4 |
| $a_{p,\mathrm{sat}}^{\mathrm{PE}}$ | Å$^2$ | 71.6 | 75.6 | 0.267 | 0.233 | 2.37 | 2.26 | 5 |
| $V_{\mathrm{HC}}^{\mathrm{DOPE}}$ | Å$^3$ | 835 | 875 | 0.255 | 0.253 | 2.19 | 2.18 | 6 |
| $\Delta V_{\mathrm{ether}}$ | Å$^3$ | -20 | -10 | 0.113 | 0.113 | 1.08 | 1.08 | 7 |
| $\alpha_V$ | K$^{-1}$ | $8\times10^{-4}$ | 0.0012 | 0.107 | 0.109 | 0.968 | 0.964 | 8 |
| $V_{\mathrm{TS}}$ | Å$^3$ | 630 | 710 | 0.0843 | 0.0431 | 0.729 | 0.371 | 9 |
| $V_{\mathrm{CH_2}}$ | Å$^3$ | 26.5 | 28.7 | 0.0258 | 0.0258 | 0.217 | 0.221 | 10 |
| $V_{\mathrm{CH=CH}}$ | Å$^3$ | 39.2 | 43.2 | 0.0254 | 0.0254 | 0.218 | 0.218 | 11 |
| $\Delta a_p^{\mathrm{PC}}$ | Å$^2$ | 1 | 3 | 0 | 0 | 0 | 0 | 12 |

**Table S3: Pairwise interaction sensitivity analysis of literature-derived input parameters to the lipid volume model.** Synergistic effects are considered for the top 4 most impactful input parameters (on average) in the univariate sensitivity analysis. Input parameters are defined in Methods. $|I|$ denotes the absolute non-additive interaction magnitude, computed as the deviation of the pairwise perturbation from the sum of the corresponding one-at-a-time perturbations. LL, LH, HL, and HH represent the combinations of low- and high-end estimates for parameters A and B. All pairwise interactions are small relative to the individual-parameter effects in Table S2, indicating that the curvature and thickness estimates are not strongly sensitive to combined parameter uncertainty.

**(A)** $c$ **of unhosted samples at 75 °C**

| Input parameter pair | A units | A low (L) | A high (H) | B units | B low (L) | B high (H) | LL $\lvert I\rvert$ (Å$^{-1}$) | LH $\lvert I\rvert$ (Å$^{-1}$) | HL $\lvert I\rvert$ (Å$^{-1}$) | HH $\lvert I\rvert$ (Å$^{-1}$) | Max $\lvert I\rvert$ (Å$^{-1}$) | Impact rank |
|---|---|---|---|---|---|---|---|---|---|---|---|---|

| | | | | | | | | | | | | |
|---|---|---|---|---|---|---|---|---|---|---|---|---|
| $R_{\text{aniso}}$ + $\kappa_V$ | | 2 | 4 | bar$^{-1}$ | 3.5×10$^{-5}$ | 5.5×10$^{-5}$ | 3×10$^{-4}$ | 3.69×10$^{-4}$ | 4.59×10$^{-4}$ | 7.81×10$^{-4}$ | 7.81×10$^{-4}$ | 1 |
| $R_{\text{aniso}}$ + $a_{p,\text{sat}}^{\text{PE}}$ | | 2 | 4 | Å$^2$ | 71.6 | 75.6 | 3.99×10$^{-5}$ | 4.04×10$^{-5}$ | 3.86×10$^{-5}$ | 3.62×10$^{-5}$ | 4.04×10$^{-5}$ | 2 |
| $R_{\text{aniso}}$ + $V_{\text{HC}}^{\text{DOPE}}$ | | 2 | 4 | Å$^3$ | 835 | 875 | 3.07×10$^{-5}$ | 3.13×10$^{-5}$ | 3.97×10$^{-5}$ | 3.99×10$^{-5}$ | 3.99×10$^{-5}$ | 3 |
| $\kappa_V$ + $a_{p,\text{sat}}^{\text{PE}}$ | bar$^{-1}$ | 3.5×10$^{-5}$ | 5.5×10$^{-5}$ | Å$^2$ | 71.6 | 75.6 | 2.01×10$^{-5}$ | 2.09×10$^{-5}$ | 1.72×10$^{-5}$ | 1.62×10$^{-5}$ | 2.09×10$^{-5}$ | 4 |
| $\kappa_V$ + $V_{\text{HC}}^{\text{DOPE}}$ | bar$^{-1}$ | 3.5×10$^{-5}$ | 5.5×10$^{-5}$ | Å$^3$ | 835 | 875 | 1.32×10$^{-5}$ | 1.33×10$^{-5}$ | 1.95×10$^{-5}$ | 1.96×10$^{-5}$ | 1.96×10$^{-5}$ | 5 |
| $a_{p,\text{sat}}^{\text{PE}}$ + $V_{\text{HC}}^{\text{DOPE}}$ | Å$^2$ | 71.6 | 75.6 | Å$^3$ | 835 | 875 | 1.07×10$^{-5}$ | 1.1×10$^{-5}$ | 9.79×10$^{-6}$ | 1×10$^{-5}$ | 1.1×10$^{-5}$ | 6 |

### (B) $c_0$ of guest lipids at 35 °C

| Input parameter pair | A units | A low (L) | A high (H) | B units | B low (L) | B high (H) | LL $\lvert I \rvert$ (Å$^{-1}$) | LH $\lvert I \rvert$ (Å$^{-1}$) | HL $\lvert I \rvert$ (Å$^{-1}$) | HH $\lvert I \rvert$ (Å$^{-1}$) | Max $\lvert I \rvert$ (Å$^{-1}$) | Impact rank |
|---|---|---|---|---|---|---|---|---|---|---|---|---|
| $R_{\text{aniso}}$ + $\kappa_V$ | | 2 | 4 | bar$^{-1}$ | 3.5×10$^{-5}$ | 5.5×10$^{-5}$ | 6.3×10$^{-5}$ | 1.07×10$^{-4}$ | 1.47×10$^{-4}$ | 2.59×10$^{-4}$ | 2.59×10$^{-4}$ | 1 |
| $a_{p,\text{sat}}^{\text{PE}}$ + $V_{\text{HC}}^{\text{DOPE}}$ | Å$^2$ | 71.6 | 75.6 | Å$^3$ | 835 | 875 | 2.49×10$^{-5}$ | 3.37×10$^{-5}$ | 1.89×10$^{-5}$ | 1.93×10$^{-5}$ | 3.37×10$^{-5}$ | 2 |
| $R_{\text{aniso}}$ + $a_{p,\text{sat}}^{\text{PE}}$ | | 2 | 4 | Å$^2$ | 71.6 | 75.6 | 1.57×10$^{-5}$ | 2.17×10$^{-5}$ | 2.94×10$^{-5}$ | 1.27×10$^{-5}$ | 2.94×10$^{-5}$ | 3 |
| $R_{\text{aniso}}$ + $V_{\text{HC}}^{\text{DOPE}}$ | | 2 | 4 | Å$^3$ | 835 | 875 | 1.03×10$^{-5}$ | 7.55×10$^{-6}$ | 1.16×10$^{-5}$ | 1.93×10$^{-5}$ | 1.93×10$^{-5}$ | 4 |
| $\kappa_V$ + $V_{\text{HC}}^{\text{DOPE}}$ | bar$^{-1}$ | 3.5×10$^{-5}$ | 5.5×10$^{-5}$ | Å$^3$ | 835 | 875 | 9×10$^{-6}$ | 6.73×10$^{-6}$ | 7.55×10$^{-6}$ | 5.55×10$^{-6}$ | 9×10$^{-6}$ | 5 |
| $\kappa_V$ + $a_{p,\text{sat}}^{\text{PE}}$ | bar$^{-1}$ | 3.5×10$^{-5}$ | 5.5×10$^{-5}$ | Å$^2$ | 71.6 | 75.6 | 7.42×10$^{-6}$ | 7.34×10$^{-6}$ | 4.81×10$^{-6}$ | 8.9×10$^{-6}$ | 8.9×10$^{-6}$ | 6 |

### (C) $h_{\text{eff}}$ of unhosted samples at 75 °C

| Input parameter pair | A units | A low (L) | A high (H) | B units | B low (L) | B high (H) | LL $\lvert I \rvert$ (Å) | LH $\lvert I \rvert$ (Å) | HL $\lvert I \rvert$ (Å) | HH $\lvert I \rvert$ (Å) | Max $\lvert I \rvert$ (Å) | Impact rank |
|---|---|---|---|---|---|---|---|---|---|---|---|---|
| $R_{\text{aniso}}$ + $\kappa_V$ | | 2 | 4 | bar$^{-1}$ | 3.5×10$^{-5}$ | 5.5×10$^{-5}$ | 0.38 | 0.548 | 0.72 | 0.963 | 0.963 | 1 |
| $R_{\text{aniso}}$ + $a_{p,\text{sat}}^{\text{PE}}$ | | 2 | 4 | Å$^2$ | 71.6 | 75.6 | 0.0552 | 0.046 | 0.0556 | 0.0655 | 0.0655 | 2 |
| $\kappa_V$ + $a_{p,\text{sat}}^{\text{PE}}$ | bar$^{-1}$ | 3.5×10$^{-5}$ | 5.5×10$^{-5}$ | Å$^2$ | 71.6 | 75.6 | 0.0366 | 0.022 | 0.0414 | 0.049 | 0.049 | 3 |
| $R_{\text{aniso}}$ + $V_{\text{HC}}^{\text{DOPE}}$ | | 2 | 4 | Å$^3$ | 835 | 875 | 0.0245 | 0.0242 | 0.0202 | 0.0194 | 0.0245 | 4 |

| | | | | | | | | | | | | |
|---|---|---|---|---|---|---|---|---|---|---|---|---|
| $\kappa_V$ + $V_{\mathrm{HC}}^{\mathrm{DOPE}}$ | $\mathrm{bar}^{-1}$ | $3.5\times10^{-5}$ | $5.5\times10^{-5}$ | $Å^3$ | 835 | 875 | 0.0121 | 0.0119 | 0.00919 | 0.00897 | 0.0121 | 5 |
| $a_{p,\mathrm{sat}}^{\mathrm{PE}}$ + $V_{\mathrm{HC}}^{\mathrm{DOPE}}$ | $Å^2$ | 71.6 | 75.6 | $Å^3$ | 835 | 875 | 0.00431 | 0.00462 | 0.00552 | 0.00549 | 0.00552 | 6 |

**Table S4: Model specifications, fitted parameters, and predictions**

**(A)** Fitted pressure model summaries for unhosted $H_{II}$ systems at 75 °C.

| Response variable | | |
|---|---|---|
| $c$ | Model formula | $c = a_j\{lipid: ts = j\} + \beta P$ |
| | Pseudo-AICc | -8186.8 |
| | RMSE | $6.03\times10^{-4}$ |
| | Weighted RMSE | $6.41\times10^{-4}$ |
| | Weighted RSS | $2.85\times10^{-4}$ |
| | N | 689 |
| $h_{\mathrm{eff}}$ | Model formula | $\log(h_{\mathrm{eff}}) = \alpha c(\mathrm{lipid, ts}, P) + \beta P + \gamma P^2$ |
| | Pseudo-AICc | -2901.4 |
| | RMSE | 0.0292 |
| | Weighted RMSE | 0.0297 |
| | Weighted RSS | 0.616 |
| | N | 700 |

**(B)** Curvature model coefficients for unhosted $H_{II}$ systems at 75 °C. Since this model is linear with no intercept, its 0-bar curvature estimates for each lipid system are identical to the system-specific coefficients. Curvatures with 12% w/w TS represent $c_0$.

| Term type | Lipid system | Coefficient | Coeff. SE | Coeff. unit |
|---|---|---|---|---|
| System-specific | diacyl POPE, 0% w/w TS | -0.037424 | $7.4\times10^{-5}$ | $Å^{-1}$ |
| | diacyl POPE, 6% w/w TS | -0.035516 | $7.8\times10^{-5}$ | |
| | diacyl POPE, 12% w/w TS | -0.035028 | $6.2\times10^{-5}$ | |
| | monoalkyl POPE, 0% w/w TS | -0.038043 | $7.5\times10^{-5}$ | |
| | monoalkyl POPE, 6% w/w TS | -0.036509 | $8.6\times10^{-5}$ | |
| | monoalkyl POPE, 12% w/w TS | -0.03791 | $7.6\times10^{-5}$ | |
| | dialkyl POPE, 0% w/w TS | -0.040744 | $7.6\times10^{-5}$ | |
| | dialkyl POPE, 6% w/w TS | -0.040433 | $8\times10^{-5}$ | |
| | dialkyl POPE, 12% w/w TS | -0.040296 | $6.2\times10^{-5}$ | |
| | diacyl SOPE, 0% w/w TS | -0.037137 | $7.4\times10^{-5}$ | |
| | diacyl SOPE, 6% w/w TS | -0.035565 | $7.6\times10^{-5}$ | |
| | diacyl SOPE, 12% w/w TS | -0.035424 | $8\times10^{-5}$ | |
| | plasmenyl SOPE, 0% w/w TS | -0.041044 | $7.7\times10^{-5}$ | |
| | plasmenyl SOPE, 6% w/w TS | -0.039847 | $8.4\times10^{-5}$ | |
| | plasmenyl SOPE, 12% w/w TS | -0.040765 | $7.6\times10^{-5}$ | |
| Common term | Pressure slope | $3.823\times10^{-6}$ | $3.7\times10^{-8}$ | $Å^{-1}$ $\mathrm{bar}^{-1}$ |

**(C)** Effective hydrocarbon thickness ($h_{\mathrm{eff}}$) model coefficients and 0-bar estimates. For the $h_{\mathrm{eff}}$ model, each system-specific coefficient is the natural logarithm of the 0-bar estimate.

| Term type | Lipid system | Coefficient | Coeff. SE | Coeff. unit | Estimate at 0 bar | Estimate SE | Estimate unit |
|---|---|---|---|---|---|---|---|
| System-specific | diacyl POPE, 0% w/w TS | 2.29414 | $2.9\times10^{-4}$ | log(Å) | 9.9159 | 0.0029 | Å |
| | diacyl POPE, 6% w/w TS | 2.2441 | $3.3\times10^{-4}$ | | 9.4319 | 0.0031 | |
| | diacyl POPE, 12% w/w TS | 2.18421 | $2.5\times10^{-4}$ | | 8.8837 | 0.0022 | |
| | monoalkyl POPE, 0% w/w TS | 2.27576 | $3\times10^{-4}$ | | 9.7353 | 0.0029 | |
| | monoalkyl POPE, 6% w/w TS | 2.22168 | $3.3\times10^{-4}$ | | 9.2228 | 0.0031 | |
| | monoalkyl POPE, 12% w/w TS | 2.15688 | $3.2\times10^{-4}$ | | 8.6442 | 0.0028 | |
| | dialkyl POPE, 0% w/w TS | 2.24935 | $3.2\times10^{-4}$ | | 9.4816 | 0.003 | |
| | dialkyl POPE, 6% w/w TS | 2.1898 | $3.3\times10^{-4}$ | | 8.9334 | 0.003 | |
| | dialkyl POPE, 12% w/w TS | 2.13162 | $2.6\times10^{-4}$ | | 8.4285 | 0.0022 | |

| | | | | | | | |
|---|---|---|---|---|---|---|---|
| | diacyl SOPE, 0% w/w TS | 2.3518 | $2.9\times10^{-4}$ | | 10.5044 | 0.0031 | |
| | diacyl SOPE, 6% w/w TS | 2.2996 | $3.2\times10^{-4}$ | | 9.9702 | 0.0032 | |
| | diacyl SOPE, 12% w/w TS | 2.23983 | $3.3\times10^{-4}$ | | 9.3917 | 0.0031 | |
| | plasmenyl SOPE, 0% w/w TS | 2.32157 | $3.2\times10^{-4}$ | | 10.1916 | 0.0033 | |
| | plasmenyl SOPE, 6% w/w TS | 2.26541 | $3.4\times10^{-4}$ | | 9.6351 | 0.0032 | |
| | plasmenyl SOPE, 12% w/w TS | 2.20539 | $3.2\times10^{-4}$ | | 9.0738 | 0.0029 | |
| Common term | Pressure quadratic term | $7.35\times10^{-9}$ | $1.8\times10^{-10}$ | log(Å) bar$^{-2}$ | – | – | – |
| | Pressure slope | $7.928\times10^{-5}$ | $3.7\times10^{-7}$ | log(Å) bar$^{-1}$ | – | – | – |

**(D)** Hosted guest-lipid $c_0$ estimates at 35°C

| Lipid system | Estimate at 0 bar | Estimate SE | Unit |
|---|---|---|---|
| diacyl POPE, 12% w/w TS | -0.02213 | $6.7\times10^{-4}$ | Å$^{-1}$ |
| monoalkyl POPE, 12% w/w TS | -0.02392 | $3.2\times10^{-4}$ | |
| dialkyl POPE, 12% w/w TS | -0.03073 | $4.6\times10^{-4}$ | |
| diacyl SOPE, 11% w/w TS | -0.02308 | $3.1\times10^{-4}$ | |
| plasmenyl SOPE, 11% w/w TS | -0.02645 | $8.2\times10^{-4}$ | |
| diacyl POPC, 12% w/w TS | 0.01197 | $2.1\times10^{-4}$ | |
| monoalkyl POPC, 12% w/w TS | 0.00617 | $3.6\times10^{-4}$ | |
| diacyl SOPC, 12% w/w TS | $-7.5\times10^{-4}$ | $3.8\times10^{-4}$ | |
| plasmenyl SOPC, 12% w/w TS | -0.00489 | $2.4\times10^{-4}$ | |

**Table S5: Pairwise z-tests comparing mechanical and membrane-dynamical properties across backbone linkage types and temperatures.** Holm familywise correction was applied separately within each variable block. *** denotes P < $2.2\times10^{-16}$.

**(A)** Experimentally determined values

| Variable | Lipid/temp. A | Lipid/temp. B | Value A | SE A | Value B | SE B | z | P | $P_{\text{adj}}$ |
|---|---|---|---|---|---|---|---|---|---|
| $-c_0$ **(Å$^{-1}$)** | monoalkyl POPE 35°C | monoalkyl POPE 75°C | 0.02392 | $3.2\times10^{-4}$ | 0.03791 | $7.6\times10^{-5}$ | -42.573 | *** | *** |
| | monoalkyl POPE 35°C | dialkyl POPE 35°C | 0.02392 | $3.2\times10^{-4}$ | 0.03073 | $4.6\times10^{-4}$ | -12.210 | *** | *** |
| | monoalkyl POPE 35°C | dialkyl POPE 75°C | 0.02392 | $3.2\times10^{-4}$ | 0.040295 | $6.2\times10^{-5}$ | -50.290 | *** | *** |
| | monoalkyl POPE 75°C | dialkyl POPE 35°C | 0.03791 | $7.6\times10^{-5}$ | 0.03073 | $4.6\times10^{-4}$ | 15.483 | *** | *** |
| | monoalkyl POPE 75°C | dialkyl POPE 75°C | 0.03791 | $7.6\times10^{-5}$ | 0.040295 | $6.2\times10^{-5}$ | -24.306 | *** | *** |
| | dialkyl POPE 35°C | dialkyl POPE 75°C | 0.03073 | $4.6\times10^{-4}$ | 0.040295 | $6.2\times10^{-5}$ | -20.724 | *** | *** |
| | diacyl SOPE 35°C | monoalkyl POPE 35°C | 0.02308 | $3.1\times10^{-4}$ | 0.02392 | $3.2\times10^{-4}$ | -1.897 | 0.0579 | 0.0579 |
| | diacyl SOPE 35°C | monoalkyl POPE 75°C | 0.02308 | $3.1\times10^{-4}$ | 0.03791 | $7.6\times10^{-5}$ | -46.791 | *** | *** |
| | diacyl SOPE 75°C | monoalkyl POPE 35°C | 0.035423 | $8\times10^{-5}$ | 0.02392 | $3.2\times10^{-4}$ | 34.915 | *** | *** |
| | diacyl SOPE 75°C | monoalkyl POPE 75°C | 0.035423 | $8\times10^{-5}$ | 0.03791 | $7.6\times10^{-5}$ | -22.589 | *** | *** |

| | | | | | | | | | |
|---|---|---|---|---|---|---|---|---|---|
| | diacyl SOPE 35°C | dialkyl POPE 35°C | 0.02308 | $3.1\times10^{-4}$ | 0.03073 | $4.6\times10^{-4}$ | -13.887 | *** | *** |
| | diacyl SOPE 35°C | dialkyl POPE 75°C | 0.02308 | $3.1\times10^{-4}$ | 0.040295 | $6.2\times10^{-5}$ | -54.853 | *** | *** |
| | diacyl SOPE 75°C | dialkyl POPE 35°C | 0.035423 | $8\times10^{-5}$ | 0.03073 | $4.6\times10^{-4}$ | 10.104 | *** | *** |
| | diacyl SOPE 75°C | dialkyl POPE 75°C | 0.035423 | $8\times10^{-5}$ | 0.040295 | $6.2\times10^{-5}$ | -48.301 | *** | *** |
| | diacyl SOPE 35°C | diacyl SOPE 75°C | 0.02308 | $3.1\times10^{-4}$ | 0.035423 | $8\times10^{-5}$ | -38.838 | *** | *** |
| | diacyl SOPE 35°C | plasmenyl SOPE 35°C | 0.02308 | $3.1\times10^{-4}$ | 0.02645 | $8.2\times10^{-4}$ | -3.852 | $1.2\times10^{-4}$ | $2.3\times10^{-4}$ |
| | diacyl SOPE 35°C | plasmenyl SOPE 75°C | 0.02308 | $3.1\times10^{-4}$ | 0.040764 | $7.6\times10^{-5}$ | -55.801 | *** | *** |
| | diacyl SOPE 75°C | plasmenyl SOPE 35°C | 0.035423 | $8\times10^{-5}$ | 0.02645 | $8.2\times10^{-4}$ | 10.916 | *** | *** |
| | diacyl SOPE 75°C | plasmenyl SOPE 75°C | 0.035423 | $8\times10^{-5}$ | 0.040764 | $7.6\times10^{-5}$ | -48.521 | *** | *** |
| | plasmenyl SOPE 35°C | plasmenyl SOPE 75°C | 0.02645 | $8.2\times10^{-4}$ | 0.040764 | $7.6\times10^{-5}$ | -17.421 | *** | *** |
| | diacyl POPC 35°C | monoalkyl POPC 35°C | -0.01197 | $2.1\times10^{-4}$ | -0.00617 | $3.6\times10^{-4}$ | -14.084 | *** | *** |
| | diacyl SOPC 35°C | plasmenyl SOPC 35°C | $7.5\times10^{-4}$ | $3.8\times10^{-4}$ | 0.00489 | $2.4\times10^{-4}$ | -9.178 | *** | *** |
| $h_{\mathrm{eff}}$ **(Å)** | monoalkyl POPE 75°C | dialkyl POPE 75°C | 8.6442 | 0.0027 | 8.4286 | 0.0022 | 61.428 | *** | *** |
| | diacyl SOPE 75°C | monoalkyl POPE 75°C | 9.3918 | 0.0031 | 8.6442 | 0.0027 | 181.941 | *** | *** |
| | diacyl SOPE 75°C | dialkyl POPE 75°C | 9.3918 | 0.0031 | 8.4286 | 0.0022 | 256.543 | *** | *** |
| | diacyl SOPE 75°C | plasmenyl SOPE 75°C | 9.3918 | 0.0031 | 9.0739 | 0.0029 | 75.663 | *** | *** |
| $k_c$ **($k_BT$)** | diacyl POPE 75°C | monoalkyl POPE 75°C | 10.324 | 0.084 | 8.416 | 0.072 | 17.280 | *** | *** |
| $K_c$ **($k_BT$)** | diacyl POPC 21°C | monoalkyl POPC 21°C | 20.751 | 1.2 | 15.707 | 0.84 | 3.437 | $5.9\times10^{-4}$ | $5.9\times10^{-4}$ |

**(B)** Values determined from MD simulations of the POPE series at 1 bar and 35 °C.

| Variable | Backbone A | Backbone B | Value A | SE A | Value B | SE B | z | P | $P_{\mathrm{adj}}$ |
|---|---|---|---|---|---|---|---|---|---|
| **APL (Å²)** | diacyl | monoalkyl | 57.424 | 0.21 | 60.768 | 0.23 | -10.784 | *** | *** |
| | diacyl | dialkyl | 57.424 | 0.21 | 58.882 | 0.20 | -5.094 | $3.5\times10^{-7}$ | $3.5\times10^{-7}$ |
| | monoalkyl | dialkyl | 60.768 | 0.23 | 58.882 | 0.20 | 6.266 | $3.7\times10^{-10}$ | $7.4\times10^{-10}$ |
| $D_T$ **(μm²/s)** large box | diacyl | monoalkyl | 7.4507 | 0.16 | 7.6662 | 0.17 | -0.935 | 0.35 | 0.35 |
| | diacyl | dialkyl | 7.4507 | 0.16 | 6.9159 | 0.14 | 2.500 | 0.0124 | 0.0247 |
| | monoalkyl | dialkyl | 7.6662 | 0.17 | 6.9159 | 0.14 | 3.405 | $6.6\times10^{-4}$ | $2.0\times10^{-3}$ |
| | diacyl | monoalkyl | 7.5143 | 0.30 | 7.4561 | 0.39 | 0.119 | 0.906 | 0.906 |

| | | | | | | | | | |
|---|---|---|---|---|---|---|---|---|---|
| $D_T$ (μm²/s) small box | diacyl | dialkyl | 7.5143 | 0.30 | 5.7595 | 0.21 | 4.760 | $1.9\times10^{-6}$ | $5.8\times10^{-6}$ |
| | monoalkyl | dialkyl | 7.4561 | 0.39 | 5.7595 | 0.21 | 3.847 | $1.2\times10^{-4}$ | $2.4\times10^{-4}$ |
| $-c_0$ (Å$^{-1}$) | diacyl | monoalkyl | 0.015961 | $5.4\times10^{-4}$ | 0.018204 | $9.4\times10^{-4}$ | -2.068 | 0.0387 | 0.0387 |
| | diacyl | dialkyl | 0.015961 | $5.4\times10^{-4}$ | 0.027347 | 0.0015 | -7.323 | $2.4\times10^{-13}$ | $7.3\times10^{-13}$ |
| | monoalkyl | dialkyl | 0.018204 | $9.4\times10^{-4}$ | 0.027347 | 0.0015 | -5.278 | $1.3\times10^{-7}$ | $2.6\times10^{-7}$ |
| $k_c$ ($k_BT$) | diacyl | monoalkyl | 18.075 | 0.29 | 13.633 | 0.52 | 7.492 | $6.8\times10^{-14}$ | $2.0\times10^{-13}$ |
| | diacyl | dialkyl | 18.075 | 0.29 | 13.066 | 0.65 | 7.004 | $2.5\times10^{-12}$ | $5.0\times10^{-12}$ |
| | monoalkyl | dialkyl | 13.633 | 0.52 | 13.066 | 0.65 | 0.681 | 0.496 | 0.496 |

### Table S6: Experiment vs. simulation comparisons.

**(A)** Intrinsic curvature $c_0$ (Å$^{-1}$). Red text indicates plasmenyl simulation data in qualitative disagreement with experiment.

| Lipid | $c_0$ simulation (neutral plane, 35 °C) | | | | | | $c_0$ experiment (hosted, pivotal plane, 35 °C) | | | | | |
|---|---|---|---|---|---|---|---|---|---|---|---|---|
| | Mean | SEM | Δ vs diacyl | SEM | % of diacyl | SEM | Mean | SEM | Δ vs diacyl | SEM | % of diacyl | SEM |
| diacyl POPE | -0.016 | $5.4\times10^{-4}$ | 0 | 0 | 100 | 0 | -0.02213 | $6.7\times10^{-4}$ | 0 | 0 | 100 | 0 |
| monoalkyl POPE | -0.018 | $9.4\times10^{-4}$ | -0.002 | 0.0011 | 114.1 | 7.0 | -0.02392 | $3.2\times10^{-4}$ | -0.00179 | $7.4\times10^{-4}$ | 108.1 | 3.6 |
| dialkyl POPE | -0.027 | $1.5\times10^{-3}$ | -0.011 | 0.0016 | 171.3 | 10.8 | -0.03073 | $4.6\times10^{-4}$ | -0.0086 | $8.1\times10^{-4}$ | 138.9 | 4.7 |
| diacyl SOPE | -0.013 | $8.1\times10^{-4}$ | 0 | 0 | 100 | 0 | -0.02308 | $3.1\times10^{-4}$ | 0 | 0 | 100 | 0 |
| plasmenyl SOPE | -0.011 | $7.1\times10^{-4}$ | 0.002 | 0.0011 | 83.1 | 7.5 | -0.02645 | $8.2\times10^{-4}$ | -0.00337 | $8.7\times10^{-4}$ | 114.6 | 3.9 |

**(B)** Monolayer effective hydrocarbon thickness $h_{\mathrm{eff}}$

| Lipid | $h_{\mathrm{eff}}$ simulation (neutral plane, 35 °C) | | | | | | $h_{\mathrm{eff}}$ experiment (pivotal plane, 75 °C) | | | | | |
|---|---|---|---|---|---|---|---|---|---|---|---|---|
| | Mean | SEM | Δ vs diacyl | SEM | % of diacyl | SEM | Mean | SEM | Δ vs diacyl | SEM | % of diacyl | SEM |
| diacyl POPE | 12.38 | 0.23 | 0 | 0 | 100 | 0 | 8.8837 | 0.0022 | 0 | 0 | 100 | 0 |
| monoalkyl POPE | 12.59 | 0.46 | 0.2 | 0.52 | 101.6 | 4.2 | 8.6442 | 0.0027 | -0.2395 | 0.0035 | 97.304 | 0.039 |
| dialkyl POPE | 12.037 | 0.06 | -0.35 | 0.24 | 97.2 | 1.9 | 8.4286 | 0.0022 | -0.4551 | 0.0031 | 94.877 | 0.034 |
| diacyl SOPE | 14.1 | 0.29 | 0 | 0 | 100 | 0 | 9.3918 | 0.0031 | 0 | 0 | 100 | 0 |
| plasmenyl SOPE | 13.79 | 0.16 | -0.3 | 0.33 | 97.8 | 2.3 | 9.0739 | 0.0029 | -0.3179 | 0.0042 | 96.615 | 0.044 |

**(C)** Monolayer bending modulus $k_c$. ND = not determined.

| Lipid | $k_c$ simulation (neutral plane, 35 °C) | | | | | | $k_c$ experiment (pivotal plane, 75 °C) | | | | | |
|---|---|---|---|---|---|---|---|---|---|---|---|---|
| | Mean | SEM | Δ vs diacyl | SEM | % of diacyl | SEM | Mean | SEM | Δ vs diacyl | SEM | % of diacyl | SEM |
| diacyl POPE | 18.08 | 0.29 | 0 | 0 | 100 | 0 | 10.324 | 0.084 | 0 | 0 | 100 | 0 |
| monoalkyl POPE | 13.63 | 0.52 | -4.44 | 0.59 | 75.4 | 3.1 | 8.416 | 0.072 | -1.91 | 0.11 | 81.52 | 0.96 |
| dialkyl POPE | 13.07 | 0.65 | -5.01 | 0.72 | 72.3 | 3.8 | ND | ND | ND | ND | ND | ND |
| diacyl SOPE | 21.8 | 1.1 | 0 | 0 | 100 | 0 | ND | ND | ND | ND | ND | ND |

| | | | | | | | | | | | |
|---|---|---|---|---|---|---|---|---|---|---|---|
| plasmenyl SOPE | 17.98 | 0.81 | -3.8 | 1.3 | 82.4 | 5.5 | ND | ND | ND | ND | ND | ND |
